# Empirical Evaluation of Single-Cell Foundation Models for Predicting Cancer Outcomes

**DOI:** 10.1101/2025.10.31.685892

**Authors:** Ahmed Roman, Shreya Johri, Riccardo Conci, Eliezer M. Van Allen, Haitham Elmarakeby

**Author notes:** Corresponding Authors: Haitham Elmarakeby, Eliezer M. Van Allen.

## Abstract

Foundation models pretrained on large-scale single-cell RNA sequencing data present a promising opportunity to advance translational cancer research. However, their utility in clinically relevant, patient-level single-cell applications remains underexplored. Here, we developed an agentic strategy to systematically evaluate twelve emerging single-cell foundation models (scFMs) and three alternative baseline approaches across seven cancer-specific tasks, including cell-type annotation, cancer subtype classification, and treatment response prediction. We assessed model performance under zero-shot, continual training, and fine-tuning conditions, conducting 1,530 supervised model-fitting runs and 200 unsupervised subsample evaluations. We found that while current scFMs excelled at certain analysis tasks, such as tumor microenvironment cell annotation, they offered limited advantages in predicting clinical and biological outcomes of cancer patients compared to simpler baseline models. These insights highlight the critical role of scFM evaluation on biologically and clinically relevant tasks for precision oncology. Beyond identifying current limitations, this assessment reveals principles that can guide future methodological innovation and the use of expanded cancer single-cell cohorts to build more biologically informed and translationally effective scFMs. The resulting agentic framework supports the autonomous discovery of emerging scFMs and facilitates their standardized integration and evaluation across cancer-related tasks.

## Introduction

Single-cell transcriptomic analysis of tumor tissues offers high-resolution insights into the cellular basis of cancer progression, therapeutic response, and microenvironmental dynamics ^1–3^. Inspired by advancements in natural language processing, foundation models pretrained on extensive single-cell datasets are increasingly applied to decode this complexity in non-cancer settings, and they may hold promise for interpreting cancer single cell data as well^4–7^. These models aim to internalize universal biological principles as "prior knowledge" ^8^, potentially improving tasks ranging from automated cell annotation to biomarker discovery. Initiatives such as CELLxGENE have facilitated these efforts by collecting and har^5–7,9^monizing the requisite large-scale data ^10^, enabling foundation models to represent cellular profiles analogously to how language models capture structure and meaning in natural language ^5–7,9^.

Despite their potential, an essential question is whether the embedded prior knowledge in these pretrained models translates into demonstrable advantages for cancer-specific problems. While existing evaluations have focused on measuring foundation models’ performance in predicting cell-level outcomes ^11–16^, whether these models can meaningfully extend from cell-level patterns to patient-level clinical or biological states that are crucial for translational impact (e.g., prognostication, therapy resistance) has not yet been thoroughly tested. This gap is especially pronounced in the cancer domain, where transcriptional heterogeneity and microenvironmental context play pivotal roles in shaping disease progression and treatment response. Yet, evaluating the utility of foundation models in elucidating the transcriptional basis of clinical and biological states in cancer is essential for advancing precision oncology and guiding therapeutic strategies. Moreover, understanding the factors that influence foundation model performance will not only help researchers select optimal analytical strategies, but also illuminate principles that can guide the design of next-generation models capable of deeper biological reasoning and broader translational impact.

Here, we present an extensive and systematic analysis, with companion agentic platform to automate updates, of the performance of leading scFMs pretrained on large-scale datasets relevant to cancer research (Figure 1). Beyond tumor microenvironment cell type annotation, our evaluation compared scFMs with multiple baseline approaches across many additional translational oncology tasks, using a variety of strategies and paradigms. In total, the supervised evaluation comprised 306 unique model–task–strategy experiments, each evaluated using five-fold cross-validation, yielding 1,530 fitted model runs. The unsupervised evaluation comprised 20 unique model–dataset experiments, each assessed across 10 independently subsampled datasets, yielding 200 evaluation runs.. Together, these analyses benchmarked the tested scFM models against three baseline approaches, providing a thorough assessment of their capabilities. Overall, scFMs did not consistently outperform conventional baselines, although cancer-specific pretraining and task-specific adaptation through continual training improved performance in selected settings, highlighting the importance of domain adaptation and rigorous task-specific benchmarking.

**Figure 1.**
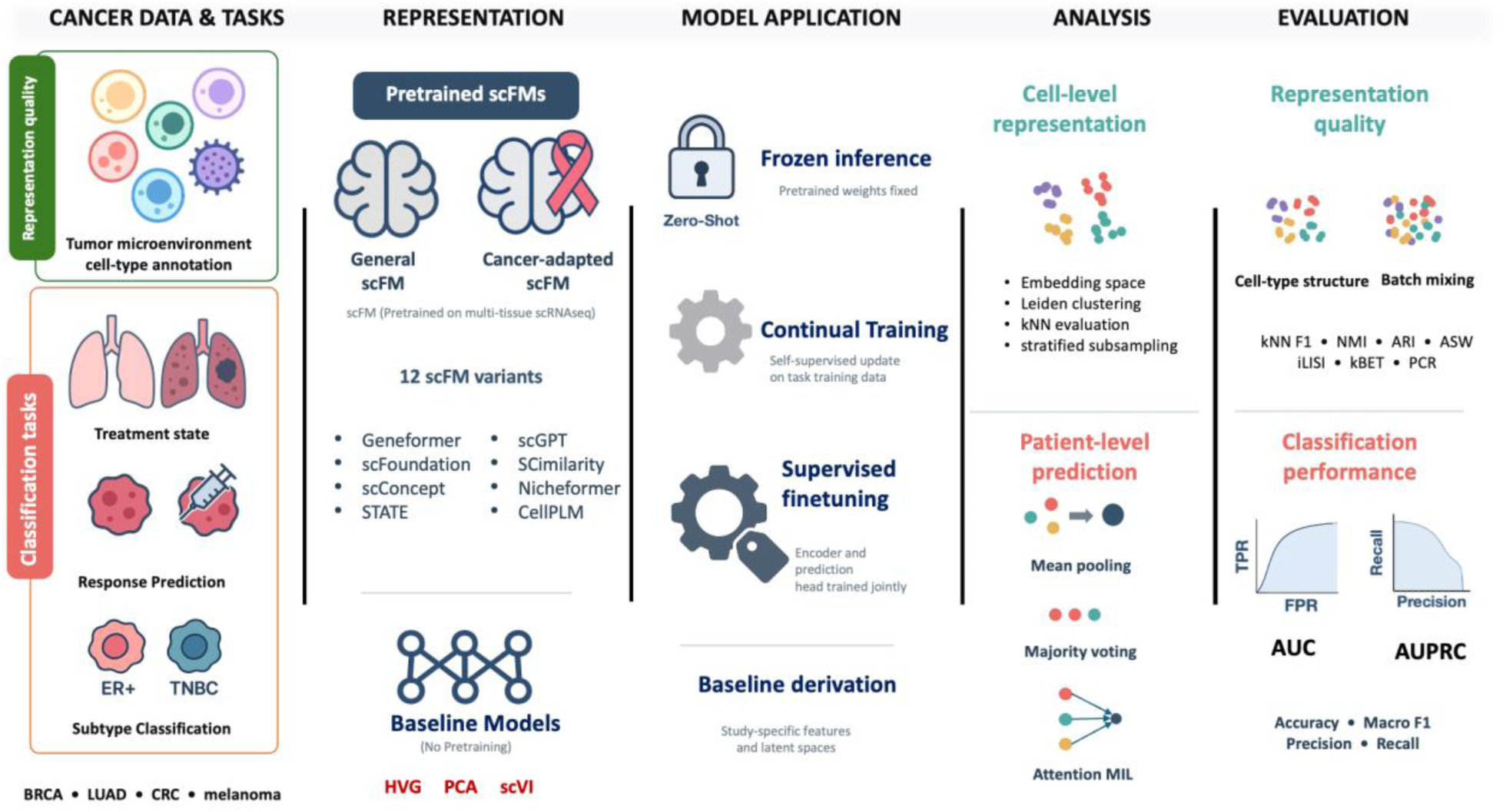
Overview of the scFM BenchScope evaluation framework. The framework evaluates 12 single-cell foundation model (scFM) variants, including general and cancer-adapted models, alongside three reference representations (HVG, PCA, and scVI). Models are assessed across seven supervised cancer-related tasks spanning cancer subtype, stage, therapeutic response, and tissue or cell-type prediction. Pretrained scFMs are evaluated using frozen inference, while selected models additionally undergo continual self-supervised pretraining or supervised fine-tuning. Cell-level representations are assessed using clustering, biological conservation, and batch-mixing metrics. For patient- or sample-level prediction, cell embeddings are aggregated using mean pooling, majority voting, or attention-based multiple-instance learning. Classification performance is evaluated using AUC, AUPRC, accuracy, macro F1, precision, and recall under matched task-specific partitions and group-aware cross-validation. The framework is accompanied by open-source software, processed evaluation data, and an interactive results dashboard.

## Results

### Single-cell embeddings show distinct trade-offs between cell-type separability and batch mixing

We first evaluated the capacity of single-cell foundation models (scFMs) to generate embeddings that capture biologically relevant structure in a zero-shot manner, without task-specific adaptation of the embedding models (see Table 1 and Methods). Using a tumor microenvironment cell annotation task with a cohort of profiled solid tumors (n = 31 tumors and ∼41K cells) and ground truth labels ^17^, we assessed the cell-type separability by training a distance-weighted k-nearest neighbor classifier (k=15) on each model’s embedding and reporting the macro F1 score on a held-out 20% test set. Training-test partitioning was performed at the cell level and stratified by cell type; consequently, cells from the same tumor could occur in both partitions. This analysis therefore measures local cell-type separability rather than generalization to unseen tumors (Figure 2A; Supplementary Figures 1–8). Across 10 stratified bootstrap subsamples, the highest cell-type separability (median kNN label F1 0.95–0.97) was reached by foundation models and simple baselines alike, with scFoundation ^7^, SCimilarity ^18^ and GF-V2-Deep^19^ ranking alongside the scVI ^20^ and PCA baselines, and CellPLM ^21^, STATE^22^, scConcept^22,23^ and GF-V2 [cancer] close behind.

**Table 1.** Architectural and pretraining characteristics of the evaluated models. Models are grouped into Geneformer, scGPT, other single-cell foundation models, and classical or non-pretrained baselines. The table summarizes the reported pretraining corpus size, parameter count, architecture, and input dimensionality of each model. “[cancer]” indicates cancer-specific pretraining, and “[finetune]” indicates task-specific fine-tuning. Classical baselines were fitted directly to each study dataset using highly variable genes (HVGs), principal component analysis (PCA), or scVI. SRT, spatially resolved transcriptomics; scRNA-seq, single-cell RNA sequencing; PCs, principal components; “dataset-dep.,” dataset-dependent; dash, not applicable or not reported.

| Model | Corpus size | Params | Depth<br>/ layers | Hidden<br>$d_{\text{model}}$ | Heads | FFN<br>$d_{\text{ff}}$ | Input<br>size |
| --- | --- | --- | --- | --- | --- | --- | --- |
| **Geneformer** |  |  |  |  |  |  |  |
| GF-V1 | Genecorpus-30M | 17M | 6 | 256 | 4 | 512 | 2048 tokens |
| GF-V1 [finetune] | Genecorpus-30M | 17M | 6 | 256 | 4 | 512 | 2048 tokens |
| GF-V2 | Genecorpus-104M | 104M | 12 | 768 | 12 | 3072 | 4096 tokens |
| GF-V2 [cancer] | Genecorpus-104M<br>+ cancer CL | 104M | 12 | 768 | 12 | 3072 | 4096 tokens |
| GF-V2 [finetune] | Genecorpus-104M | 104M | 12 | 768 | 12 | 3072 | 4096 tokens |
| GF-V2-Deep | Genecorpus-104M | 316M | 18 | 1152 | 18 | 4608 | 4096 tokens |
| **scGPT** |  |  |  |  |  |  |  |
| scGPT | 33M cells | 51M | 12 | 512 | 8 | 512 | 2048 tokens |
| scGPT [cancer] | 33M cells<br>+ 5.7M cancer | 51M | 12 | 512 | 8 | 512 | 2048 tokens |
| **Other single-cell foundation models** |  |  |  |  |  |  |  |
| scConcept | 40M cells | 30M | 8 | 512 | 8 | 1024 | 1000 tokens |
| SCimilarity | 7.9M train<br>23.4M reference | 31M | 2 encoder<br>+ 2 decoder | 1024 hidden<br>128 latent | — | — | $n_{\text{genes}}$ |
| Nicheformer | SpatialCorpus-110M | 49M | — | — | — | — | — |
| CellPLM | >9M scRNA-seq<br>+ 2M SRT | 82.4M | 4 encoder<br>+ 2 decoder | 1024 hidden<br>512 latent | — | — | 13,500 scRNA genes<br>1,000 SRT genes |
| scFoundation | >50M cells | 100M | 12 encoder<br>+ 6 decoder | 768 encoder<br>512 decoder | 12 encoder<br>8 decoder | — | 19,264 genes<br>100 bins |
| STATE | >100M perturbed<br>+ 167M observational | 600M | 8 | 512 | 16 | 1024 | 19,790 genes<br>token dim 5120 |
| **Classical / non-pretrained baselines** |  |  |  |  |  |  |  |
| HVG | study data | — | — | — | — | — | selected genes |
| PCA [20] | study data | — | — | — | — | — | 20 PCs |
| PCA [50] | study data | — | — | — | — | — | 50 PCs |
| PCA [100] | study data | — | — | — | — | — | 100 PCs |
| scVI | study data | dataset-dep. | 1 encoder<br>/ decoder NN | 128 hidden<br>10 latent | — | — | $n_{\text{genes}}$ |

**Figure 2.**
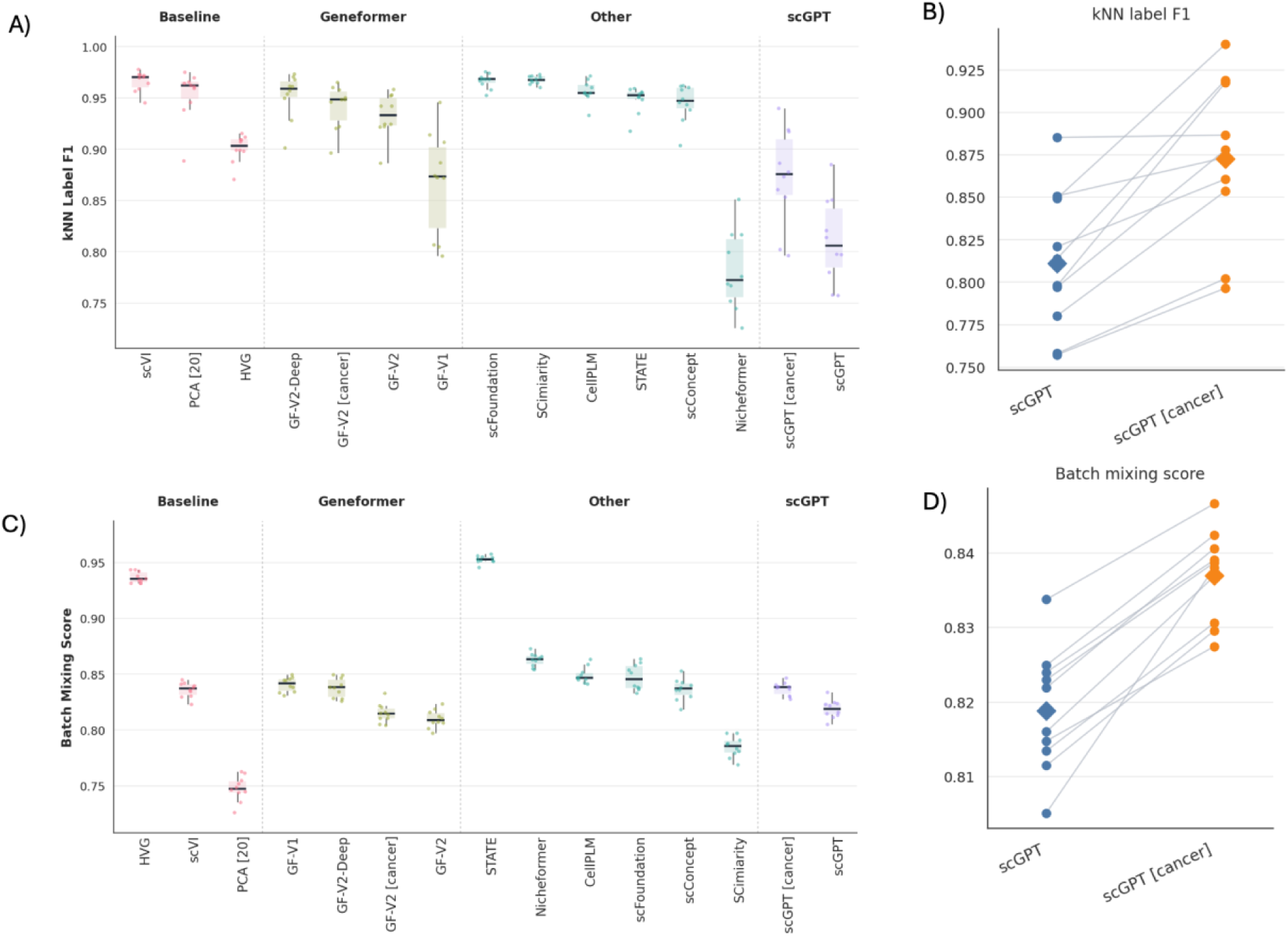
Performance of different models on annotating the tumor microenvironment. A) kNN label F1 across single-cell baseline and foundation model embeddings. Cell-type preservation was assessed by training a k-nearest-neighbor classifier (k = 15, distance-weighted) on each model’s embedding and reporting macro-averaged F1 on a held-out 20% stratified test split (higher = better cell-type separability; range 0–1). Scores are shown for 15 models across 10 stratified bootstrap subsamples (10,000 cells each). Models are grouped as Baseline, Geneformer, Other, and scGPT, and ordered within each group by median score. Box plots show the interquartile range; points are individual bootstrap runs. B) Slope plot showing that cancer-specific domain adaptation helps the scGPT model to achieve statistically significantly higher kNN label F1 (Wilcoxon P = 0.002). C) Batch mixing score across single-cell foundation model embeddings. Batch mixing was quantified to measure how well technical batches overlap within each cell type in embedding space (higher = better mixing; range 0-1).). Baseline methods (e.g., HVG) and STATE achieved among the highest scores (∼0.93–0.95), while PCA [20] and SCimilarity showed comparatively lower batch mixing (∼0.75–0.79). D) Slope plot showing that cancer-specific domain adaptation helps the scGPT model to achieve a statistically significantly higher batch mixing score (Wilcoxon P = 0.002).

We subsequently evaluated whether cancer-specific pretraining improved performance in a representative model (scGPT; Figure 2B; Methods). Across all paired subsamples, the cancer-pretrained version of scGPT (scGPT [cancer]) achieved a significant increase in kNN label F1 relative to the original, general-domain scGPT modell (Figure 2B) (Wilcoxon *P* = 0.002). Its absolute performance nevertheless remained below that of the leading baseline and foundation-model embeddings, indicating that cancer-specific pretraining can substantially improve the biological information captured by a model’s embeddings without necessarily making it competitive with alternative architectures or conventional representations.

We also evaluated batch mixing to assess the local mixing of cells from different technical batches within individual cell types in each embedding space (Figure 2C; Methods). STATE achieved the highest batch mixing score (followed by the HVG^24^ baseline), whereas PCA and SCimilarity showed comparatively limited batch integration. The Geneformer, scFoundation, CellPLM, scConcept, and scGPT embeddings showed intermediate performance. These rankings differed from those obtained using kNN label F1; SCimilarity ranked among the strongest models for cell-type classification yet showed comparatively weak batch mixing, while HVG showed strong batch mixing despite relatively low kNN label F1. STATE performed strongly on both measures, suggesting its embeddings preserved cell-type information while promoting mixing across technical batches within cell types. Cancer-specific pretraining similarly improved batch mixing: scGPT [cancer] significantly outperformed the general-domain scGPT model across the paired subsamples (Figure 2D) (Wilcoxon P = 0.002; Figure 2D), mirroring the improvement observed for kNN label F1. Similarly, the cancer-pretrained GF-V2 [cancer] also increased both metrics relative to generic GF-V2. The gain in batch mixing score was significant (Wilcoxon P = 0.006), whereas the increase in kNN label F1 was not (Wilcoxon P = 0.23; Supplementary Figure 9)

Together these results indicate that embedding quality was not determined solely by model size, pretraining-corpus scale, or membership in a particular model family. Simple baseline representations remained competitive with and in some cases outperformed larger foundation models. Moreover, the improvements associated with cancer-specific pretraining in scGPT and Geneformer suggest that domain-relevant training can improve multiple properties of a learned representation. Joint evaluation of cell-type separability and batch mixing provides a more complete assessment of embedding quality, as the two metrics capture complementary and sometimes discordant properties of single-cell embedding spaces.

### Continual training improves embedding structure and performance

We next assessed whether task-specific continual pretraining on each evaluation dataset altered embedding structure and improved quantitative performance (Figure 3). This intervention used the cells from the downstream dataset without using their task labels and was therefore distinct from supervised fine-tuning and cancer-specific pretraining. UMAP visualizations were used to qualitatively compare scVI, GF-V1 [zero-shot], and GF-V1 [continual] embeddings (Figure 3A). The zero-shot GF-V1 embeddings captured broad cell-type structure; however, continual training produced more compact cell-type regions, as well as visually clearer boundaries between several major tumor microenvironment populations. Although these visualizations are qualitative, they visually suggest that task-specific continual training can reorganize the embedding space in a way that aligns more closely with annotated cell identities.

**Figure 3.**
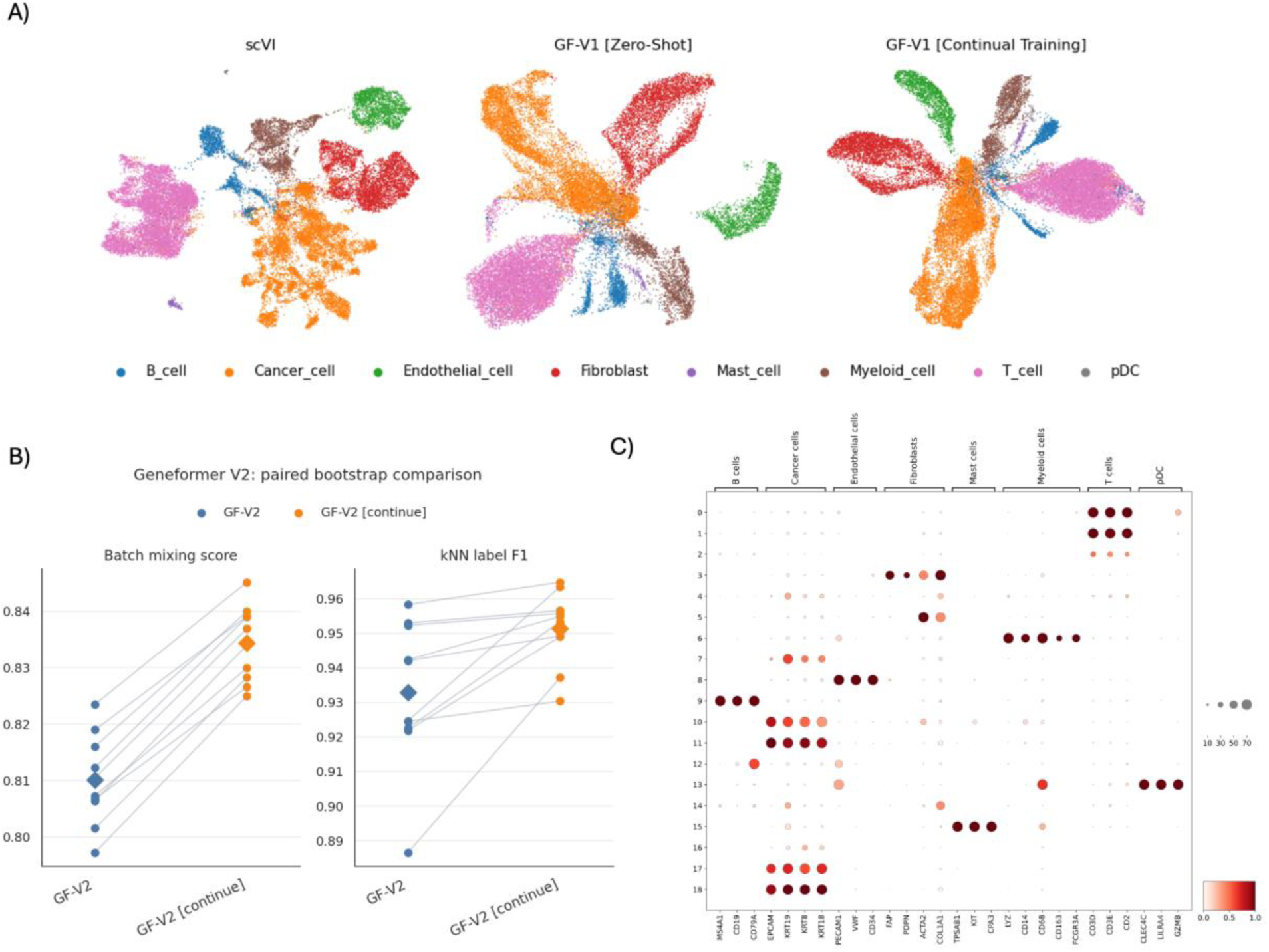
Effect of continual training on model performance. A) UMAP plots illustrating cell-type separability across representative methods: scVI (left), GF-V1 [zero-shot] (middle), and GF-V1 [continual] (right). GF-V1 embeddings after continual training show markedly improved intra-cluster compactness and more precise inter-cluster boundaries, highlighting the benefits of domain-adaptive continual pretraining. B) Slope plot showing that continual helps the GF-V2 model to achieve statistically significantly higher batch mixing score and kNN label F1. batch mixing score (Δ = +0.024; 95% CI, 0.020–0.028; Holm p = 0.004) and kNN label F1 (Δ = +0.019; 95% CI, 0.004–0.048; Holm p = 0.004). Each dot represents a subsample of (10,000 cells). C) Dot plot of canonical tumor microenvironment (TME) marker genes across Leiden clusters (resolution 0.3) derived from GF-V2 [continual] embeddings in pre-treatment BRCA tumor cells (n=26,697 cells, 19 clusters). Dot size indicates the fraction of cells with detectable expression; color indicates mean log-normalized expression (scaled per gene). Embedding-derived clusters recovered expected marker structure, with the strongest enrichments including Mast cells markers in cluster 15 (89% expressing cells), Cancer cells markers in cluster 18 (99% expressing cells), pDC markers in cluster 13 (94% expressing cells). Per-cluster marker statistics are provided in the accompanying tables.

We also quantified the effect of task-specific continual training using paired comparisons of GF-V2 and GF-V2 [continual] across 10 stratified bootstrap subsamples of 10,000 cells each (Figure 3B). The batch mixing score increased by 0.024 (95% CI, 0.020–0.028; Holm-adjusted p = 0.004), and kNN label F1 increased by 0.019 (95% CI, 0.004–0.048; Holm-adjusted p = 0.004). These paired improvements indicate that continual training boosted cell-type separability without degrading batch integration and, in this comparison, improved both properties simultaneously in this comparison.

To determine whether the structure recovered from the continually trained embeddings aligned with established tumor microenvironment biology, we examined canonical marker expression across Leiden clusters derived from the GF-V2 [continual] embeddings (Figure 3C). The resulting clusters showed distinct enrichment for expected lineage markers. Among the strongest lineage-specific patterns, 89% of cells in cluster 15 exhibited mast-cell markers, 99% of cells in cluster 18 exhibited cancer-cell markers, and 94% of cells in cluster 13 exhibited plasmacytoid dendritic-cell markers. Thus, continual training increased both kNN label F1 and batch mixing in GF-V2, while clusters derived from GF-V2 [continual] embeddings showed coherent canonical marker expression. A similar observation was shown for Geneformer GF-V1 embeddings (Supplementary Fig. 10). Together, these findings provide complementary qualitative, quantitative, and marker-based evidence that cancer-specific continual training could improve model representations of the tumor microenvironment.

### Predictive performance of scFMs in translational oncology tasks

Next, we assessed the supervised predictive performance of scFMs across a wide range of clinically relevant classification tasks derived from lung adenocarcinoma (LUAD),breast cancer (BRCA), colorectal (CRC), and melanoma cohorts. In each task, we aimed to predict patient-level clinical outcomes using aggregated information from single-cell profiles. These tasks were designed to capture distinct biological contrasts and span a spectrum of prediction difficulties, thus providing a systematic benchmarking framework. Specifically, we included: (i) distinguishing treatment-naïve from tyrosine kinase inhibitor (TKI)-treated patients using cancer cells; (ii) differentiating treatment-naïve from anti-PD1–treated BRCA patients using T-cells; (iii) classifying estrogen receptor positive (ER+) versus triple-negative breast cancer (TNBC) patients based on cancer cells; (iv) predicting immunotherapy response in melanoma patients; (v) distinguishing mismatch repair-deficient (MMRd) from mismatch repair-proficient (MMRp) tumors in CRC; (vi) differentiating exhausted versus non-exhausted T-cell states in BRCA tumors; and (vii) distinguishing treatment-naïve from neoadjuvant chemotherapy–treated BRCA patients using cancer cells. Then, we pursued a systematic evaluation of scFMs for each of these cancer outcome predictions across multiple analytical dimensions: comparing predictive performance across models, quantifying the effects of model scale and cancer-specific adaptation, comparing patient-level aggregation strategies, and assessing gains from end-to-end fine-tuning over frozen embeddings (Methods).

#### Performance across cancer classification tasks

We first examined patient-level prediction performance across seven cancer classification tasks using multiple-instance learning (MIL), with performance summarized using the mean AUPRC across five cross-validation experiments (Figure 4A; other aggregation strategies are reported in Supplementary Figure 11). In this setting, performance depended on both the embedding model and the biological context being evaluated, reflecting how well clinically relevant differences were preserved in each representation. Mean AUPRC fell within a narrow band for most models (∼0.85–0.94). SCimilarity ranked highest (∼0.94), followed by STATE (∼0.92). However, the scVI (∼0.90), PCA (∼0.89) and HVG (∼0.88) baselines were interleaved with the strongest foundation models, and only the two scGPT variants (∼0.77–0.79) fell clearly below this band. Within the Geneformer family, GF-V2 [cancer] and GF-V2 (∼0.89–0.90) both exceeded the deeper GF-V2-Deep (∼0.86) and the earlier GF-V1 (∼0.85), indicating that neither increased model depth nor cancer-specific pretraining provided a clear performance advantage for that model strategy. Per-model values for all fifteen models are shown in Figure 4A.

**Figure 4.**
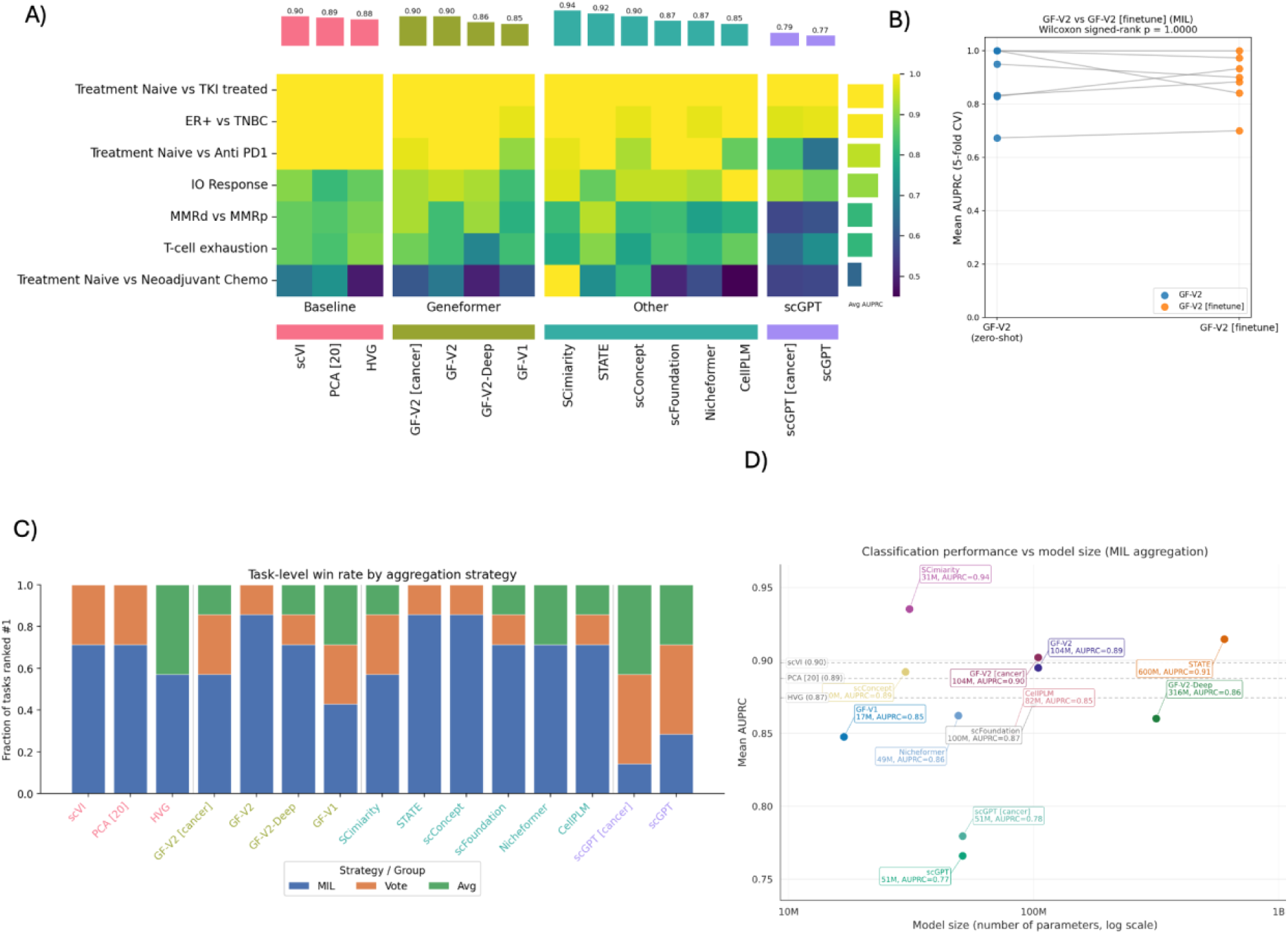
Evaluation of performance of outcome prediction. A) Comparison between all models in predicting seven different outcomes measured using the area under the precision-recall curve, AUPRC. Each cell represents the average of five cross-validation folds. The bars on top show the average performance of each model across the different tasks. B) Frozen inference (zero-shot) vs. finetuned Geneformer V2 (104M) on cancer classification benchmarks. Mean task-level AUPRC (5-fold CV, MIL) on seven cancer classification benchmarks. Finetuning showed no significant overall benefit (mean ΔAUPRC = −0.008, Wilcoxon p = 1.00). C) Task win rate across 7 classification benchmarks per model. MIL ranks first on 63% of model–task pairs; Vote (21%) and Avg (16%) are preferred for baselines and scGPT. Geneformer and most other foundation models favor MIL on most tasks. D) Classification performance versus model size. Mean area under the precision–recall curve (AUPRC) for each embedding method using MIL aggregation and 5-fold cross-validation, plotted against the number of model parameters (log scale). Each point is a foundation model (n = 12). Reference baselines (HVG, PCA, scVI) are shown as dashed horizontal lines. Performance does not increase monotonically with model size: SCimilarity (31M parameters) achieves the highest AUPRC, while several mid-sized models outperform much larger ones and simple baselines remain competitive with many pretrained embeddings.

Across the separate classification tasks, performance showed marked variation, with most models producing strong separation when distinguishing treatment-naïve LUADs from those exposed to TKI, ER+ cases from TNBC samples, or treatment-naïve tumors from anti-PD1-exposed specimens. However, other tasks resulted in heterogeneous model performances, including predicting immunotherapy response, mismatch-repair status or T-cell exhaustion. Classification of treatment-naïve cases against those after neoadjuvant chemotherapy resulted in the greatest spread in outcomes among models, with SCimilarity achieving the highest single-task AUPRC in this setting.

We also compared end-to-end fine-tuning of GF-V2 for classification with its frozen-embedding counterpart, in which Geneformer parameters remained fixed and a separate classifier was trained on the resulting embeddings (Figure 4B). On average, the fine-tuned model achieved similar AUPRCs to that of the frozen model. One possible explanation is that the limited number of patient samples was insufficient to support effective end-to-end optimization. Fine-tuning did not produce consistent improvements across tasks, as gains on some tasks were offset by declines on others, and a Wilcoxon signed-rank test showed no systematic task-level improvement for GF-V2 (mean ΔAUPRC = −0.008; p = 1.00). This lack of a consistent advantage was also reflected in the 95% bootstrap confidence interval (−0.11 to +0.04) and non-significant results for both GF-V1 [finetune] (p ≈ 0.688) and GF-V2 [finetune] (p ≈ 1.00) (Supplementary Figure 12).

Thus, improvements in embedding structure, cell-type separability, and batch mixing observed with task-specific continual pretraining did not reliably translate into improved clinical endpoint prediction. More broadly, these findings illustrate that improvements in representation quality alone do not guarantee consistent gains across heterogeneous downstream prediction tasks. The limited sizes of the evaluated patient cohorts may have contributed to this discrepancy, although the present analyses were not designed to establish its cause.

#### Comparing aggregation strategies for patient-level predictions

We next compared three aggregation strategies for generating patient-level predictions. Since all the evaluated models produce cell-level embeddings, we applied three strategies to aggregate these embeddings or their downstream predictions at the patient level: (i) multiple-instance learning (MIL) ^25,26^, (ii) majority voting^27^, and (iii) embedding averaging^25,28^ (Methods; Figure 4A,C; Supplementary Figures 13,14,15). In MIL, an attention-based model learns to assign different weights to individual cells when forming a patient-level prediction. Embedding averaging instead compresses each patient into one representation by taking the mean of that patient’s cell embeddings. Finally, in majority voting, a classifier first generates a prediction for each cell, and the most frequently predicted outcome is assigned to the patient.

Across the seven tasks, MIL achieved the highest AUPRC in 63% of model-task comparisons, followed by majority voting (21%) and embedding averaging (16%) (Figure 4C). MIL achieved the highest AUPRC in six of seven tasks for GF-V2, STATE, and scConcept, and in five of seven tasks for GF-V2-Deep, scFoundation, Nicheformer, and CellPLM. However, scGPT variants more often performed best with majority voting and embedding averaging. These results suggest that, for most tasks, patient-outcome information is encoded not only in the average or median cellular state but also in the within-patient heterogeneity and multivariate structure of single-cell representations, which MIL can preserve and exploit more effectively than aggregation by averaging or voting. Nevertheless, embedding averaging and majority voting can be advantageous when outcome-relevant information is primarily captured by population-level shifts in the mean or by the prevalence of particular cellular states, quantities that these simpler aggregation strategies can estimate robustly.

Results on the seven-task subset reinforced the coupling between aggregation strategy and representation model. Under embedding averaging, the PCA, scVI, and HVG baselines attained the highest mean AUPRCs (∼0.90), ahead of the Geneformer models and SCimilarity (0.87–0.89), and scFoundation, CellPLM, and the scGPT models (0.79–0.83). Under majority voting, the performance range narrowed (0.79–0.87), with HVG and GF-V2 achieving the highest mean AUPRCs. Notably, the scGPT family performed better under majority voting than under embedding averaging, further illustrating that the optimal aggregation strategy depended, in part, on the underlying representation model.

Overall, MIL was the aggregation method that most often produced the strongest performance, although its advantage varied across embedding models: baseline and most other foundation representations benefited from MIL, while representations of GF-V1, scGPT and scGPT [cancer] more frequently favored embedding averaging or majority voting. These findings indicate that aggregation strategy should be considered jointly with the representation model rather than treated as a universally fixed downstream component. The aggregation strategy should be treated as a design choice and optimized as a hyperparameter during model development.

#### Impact of Model Size and Cancer-Specific Adaptation on scFM Performance

Finally, we examined whether model size or cancer-specific pretraining was associated with mean AUPRC across the clinical prediction tasks. Performance did not increase monotonically with parameter count (Figure 4D). SCimilarity, despite having only 31M parameters, achieved the highest mean AUPRC (∼ 0.94), followed by STATE (∼0.92), which contains roughly 600M parameters. A similar pattern was observed within the GF-V2 family: GF-V2-Deep (316M parameters) obtained a mean AUPRC near 0.86, yet it was exceeded by the smaller GF-V2 [cancer] (∼ 0.90) and by GF-V2 (about 0.89). scConcept also performed strongly (about 0.89), while scFoundation (about 0.87), Nicheformer (about 0.86), GF-V1 (about 0.85), and CellPLM (about 0.85) showed intermediate performance.

Cancer-specific pretraining produced only modest differences within matched model families. GF-V2 [cancer] slightly outperformed GF-V2, while scGPT [cancer] (∼ 0.78) marginally outperformed scGPT (∼ 0.77), although both scGPT variants had the lowest mean AUPRC in this comparison. Notably, conventional baselines, with significantly smaller parameter count, including scVI (∼ 0.90), PCA (∼ 0.89), and HVG (∼ 0.87), remained competitive against multiple foundation models. Overall, these findings indicate that parameter count alone does not explain predictive performance in these clinical settings. Smaller models and standard baselines, frequently matched or outperformed substantially larger architectures, while cancer-specific pretraining yielded only modest improvements within the matched Geneformer and scGPT model families.

### Agentic automation of scFMs

Finally, to facilitate ongoing evaluation and benchmarking of scFMs for cancer purposes as the field evolves and new scFMs are introduced into the scientific community, we developed an agentic platform (scFM BenchScope; Methods; https://marakeby.github.io/scfm-benchscope/) that automates model discovery, model assessment, and pursues standardized, human-reviewed integration and reevaluation in the context of all prior models recovered to date.

## Discussion

In this systematic benchmark, we evaluated single-cell foundation models (scFMs) across complementary dimensions of performance, including preservation of biological structure, batch mixing, patient-level clinical prediction, model scale, cancer-specific pretraining, task-specific continual pretraining, supervised fine-tuning, and patient-level aggregation. Several major findings emerged. First, scFMs generated embeddings that captured biologically meaningful cellular structure, but they did not consistently outperform conventional representations such as PCA, highly variable genes, and scVI. Second, additional training on cancer or task-specific data improved some embedding properties, although the magnitude and direction of these effects depended on the model and metric. Third, patient-level classification performance did not increase monotonically with model size, and a relatively small model, SCimilarity, achieved the highest mean AUPRC under MIL aggregation. Fourth, end-to-end fine-tuning of Geneformer V1 and V2 did not consistently improve performance over their frozen representations under the evaluated protocols. Finally, patient-level performance depended substantially on how cell-level information was aggregated: attention-based multiple-instance learning (MIL) most frequently achieved the highest AUPRC, but simpler approaches were preferable for some representations. Together, these findings show that model scale, embedding quality, domain relevance, adaptation strategy, and aggregation method represent distinct dimensions of performance rather than interchangeable indicators of model utility.

The embedding analyses showed that high cell-type separability was not unique to foundation models. Several scFMs achieved high kNN label F1, but scVI and PCA were similarly competitive, indicating that the dominant transcriptional variation defining major tumor-microenvironment cell types can be recovered without large-scale pretraining. This finding does not imply that scFMs contain no information beyond conventional representations. Instead, it indicates that broad cell-type identity may be an insufficiently discriminating endpoint for demonstrating the added value of pretraining, particularly when the annotated populations are separated by strong transcriptional programs. Foundation models may encode more subtle cell states, regulatory programs, or relationships that are not captured by a cell-type classification probe. Conversely, strong performance on cell-type recovery should not be interpreted as evidence that an embedding contains the information required for a specific clinical prediction task.

Model rankings also differed across biological-conservation and batch-mixing metrics. Representations that strongly preserved cell-type identity were not necessarily those that most effectively mixed cells across batches, emphasizing that these objectives can capture different (and sometimes competing) properties of an embedding. In cancer data, apparent batch differences may additionally reflect differences in tissue, treatment, cellular composition, or disease state rather than technical variation alone. Consequently, maximizing batch mixing without considering biological context could remove clinically relevant heterogeneity. These results support evaluating embeddings using multiple complementary metrics rather than reducing representation quality to a single composite score.

Cancer-specific pretraining produced some of the clearest improvements at the embedding level. Relative to the general scGPT model, scGPT [cancer] improved both cell-type separability and batch mixing. GF-V2 [cancer] also improved batch mixing relative to GF-V2, although its increase in kNN label F1 was not statistically significant. The corresponding gains in patient-level classification were comparatively modest for both model families. One possible interpretation is that exposure to cancer data improves the overall organization of tumor-derived transcriptomes without necessarily enriching the representation for every clinical endpoint. Patient outcomes may depend on comparatively subtle or rare cellular states, treatment-specific processes, and patient-level compositional differences that are not directly optimized during cancer-specific pretraining. Thus, improved alignment with the cancer domain should not be assumed to confer task-specific predictive benefit.

Task-specific continual pretraining likewise had model-dependent effects. Continual pretraining improved both kNN label F1 and batch mixing for GF-V2, whereas GF-V1 showed a nonsignificant increase in cell-type separability accompanied by a small decrease in batch mixing. These contrasting results suggest that the value of continual pretraining depends on the starting checkpoint, model capacity, and compatibility between the pretrained representation and the additional training data and objective. They also argue against treating continual pretraining as a uniformly beneficial intervention. Because the continual-pretrained variants were evaluated primarily through embedding-level analyses, whether these changes improve patient-level clinical prediction remains to be established directly.

A central finding of this study is that representation quality and clinical utility cannot be inferred from one another. Cell-type separability evaluates whether established biological identities are accessible from an embedding, whereas patient-level prediction asks whether cells collectively encode information about treatment exposure, molecular phenotype, immune state, or therapeutic response. These endpoints may depend on different aspects of the data. Cell identity is often represented by large, recurrent expression programs, while clinical outcomes may depend on subtle state changes, rare populations, interactions between tumor and immune compartments, or differences in cellular composition. An embedding can therefore organize canonical cell types effectively while discarding, compressing, or failing to expose outcome-relevant variation. This distinction is important for scFM evaluation: embedding metrics are useful measures of biological representation, but they should not be treated as surrogate endpoints for clinical prediction.

Patient-level classification performance also did not increase monotonically with parameter count. Under MIL aggregation, SCimilarity achieved the highest mean AUPRC despite having substantially fewer parameters than several competing models, and the standard GF-V2 model outperformed the larger GF-V2-Deep variant. These observations demonstrate that parameter count alone is not a sufficient criterion for selecting an scFM. However, the benchmark compared complete modeling systems that differed in architecture, pretraining objective, training corpus, gene vocabulary, input representation, and embedding-extraction procedure. It therefore cannot isolate a causal effect of scale or establish that smaller models are intrinsically superior. SCimilarity’s strong performance in both cell-type separability and patient-level prediction is consistent with the possibility that its architecture and training objective produce a representation that is readily usable by downstream models, but the contribution of individual design choices could not be determined here. More broadly, the competitiveness of both smaller scFMs and conventional baselines suggests that alignment between a representation and the downstream task may be more consequential than parameter count alone.

The absence of a consistent benefit from end-to-end Geneformer fine-tuning was another notable result. Fine-tuning produced gains on some clinical tasks and losses on others, resulting in no systematic improvement over frozen Geneformer embeddings for either GF-V1 or GF-V2. Several explanations are possible. The limited number of patients may have been insufficient to estimate the large number of trainable parameters reliably; full-model optimization may have disrupted useful pretrained features; or the evaluated hyperparameters may not have been optimal for each dataset. Ceiling effects on tasks for which frozen embeddings already performed strongly may also have limited the opportunity for improvement. The present analysis was not designed to distinguish among these possibilities, and the findings should not be interpreted as evidence that fine-tuning is generally ineffective. Instead, they establish that, under the evaluated protocols, its additional computational complexity did not provide a reproducible advantage over frozen representations. Frozen embeddings with a carefully designed patient-level classifier therefore represent an important practical baseline against which more complex adaptation strategies should be compared.

How cell-level information was converted into a patient-level prediction was itself a major determinant of performance. Attention-based MIL most frequently achieved the highest AUPRC across representation–task comparisons, but its advantage was not universal. The standard scGPT model more frequently favored majority voting, whereas scGPT [cancer] more frequently favored embedding averaging or voting. These differences indicate that the utility of an aggregation method depends on the geometry and information content of the underlying representation.

One possible explanation for the frequent advantage of MIL is its ability to assign unequal weights to individual cells, allowing highly informative cells to contribute more strongly than they would under uniform averaging. This mechanism could be advantageous when outcome-associated signals are concentrated in particular cellular states or subpopulations. In contrast, embedding averaging may perform well when an outcome is reflected by a broad shift in the mean cellular representation, while majority voting may be effective when the prevalence of cells assigned to an outcome-associated state is informative. However, the implemented MIL model applies attention-weighted pooling after transforming individual cell embeddings; it does not explicitly model pairwise cell–cell interactions or higher-order distributional structure. Its superior performance therefore should not be taken as evidence that it captured cellular co-occurrence, covariance, or biological interactions. Moreover, attention weights were not validated as biological explanations. Targeted attention analyses, cell-removal experiments, and subpopulation-level ablations will be needed to determine which cellular signals underlie the observed advantage.

These findings have several implications for the development and evaluation of scFMs in translational oncology. First, model selection should be based on the intended biological or clinical endpoint rather than on model size, pretraining scale, or performance on a single embedding metric. Second, strong conventional baselines remain essential: without them, the benefit attributed to a foundation model may instead reflect preprocessing, dimensionality reduction, or downstream model design. Third, adaptation and aggregation should be treated as components of the complete predictive pipeline. In particular, aggregation strategy should be considered a model-design choice that can be optimized during development using training and validation data, while independent test data remain reserved for estimating generalization performance. Finally, clinical benchmarks should include patient-level endpoints because cell-level annotation alone cannot establish translational utility.

This study has several important limitations. First, our benchmarks focused on cancer-related single-cell datasets with relatively small numbers of patients, particularly for clinically relevant outcome tasks for which the patient, rather than the individual cell, is the effective independent sample. These limited sample sizes, typically a few tens of patients or fewer, increase the risk of downstream overfitting, widen uncertainty in performance estimates, and make small differences and model rankings difficult to interpret reliably. Larger and more heterogeneous cohorts will therefore be needed to determine whether the observed results generalize across institutions, cancer types, and treatment paradigms. In addition, although we systematically evaluated multiple aggregation strategies and domain-adaptation approaches, we did not exhaustively explore additional strategies, such as hierarchical or biologically informed methods that explicitly represent cell-state composition, rare subpopulations, cell-cell interactions, or spatial organization within each patient, and that may therefore yield more informative patient-level representations. Furthermore, neural downstream models could benefit from model-specific hyperparameter optimization, adaptive learning-rate schedules, and early stopping. These techniques were intentionally excluded to maintain a controlled comparison space, although their exclusion may affect models differently. Moreover, threshold-based metrics, including accuracy, precision, recall, and F1, were calculated using a fixed default threshold of 0.5, and thresholding can lead to different results; AUPRC, however, is threshold-independent and therefore is not affected by this limitation. Finally, all comparisons in this study are based solely on single-cell transcriptional data, without accounting for other data modalities inferable from single-cell analysis (e.g., copy number information, an essential dimension of tumor biology that captures genomic alterations driving cancer progression and treatment response).

Broadly, our work underscores the significant methodological and conceptual challenges that constrain the clinical application of single-cell foundation models (scFMs). Many current scFMs rely on gene-level input representations and training objectives that do not explicitly encode gene-regulatory relationships, pathways, or higher-order biological organization. These design choices may limit how well these models capture the multiscale structure of biological systems, particularly within heterogeneous tumors. Moreover, most scFMs are trained primarily at the individual cell level and do not explicitly model population composition, cell–cell communication, or tissue-level organization. Biological and clinical context is also represented inconsistently across the models, including tissue type, disease state, treatment exposure, and microenvironmental setting. Similarly, most models do not natively learn sample-level representations, leaving clinically relevant patient-level structure to be recovered through a separate downstream aggregation procedure. This separation may help explain why strong cell-level embedding quality did not consistently translate into superior patient-level prediction and why conventional representations remained highly competitive. Expanding dataset size alone is unlikely to overcome these limitations. Nonetheless, it remains essential to collect samples from more patients across a diverse range of cancer types and treatment settings, while ensuring that foundation models explicitly integrate the full tumor microenvironment to capture intercellular interactions and patient-level contextual biological signals. Future progress will require careful attention to clinical and biological context, improved patient-level and multiscale representation learning, and the incorporation of structured prior knowledge that reflects biological organization and constraints. Advances in loss function design and representation learning will also be necessary to develop models that can reason in biological terms and provide interpretable, mechanistically grounded insights for downstream tasks in cancer research and beyond.

## Conclusion

In this benchmarking study, we systematically evaluated single-cell foundation models (scFMs) across an extensive set of tasks, revealing several important insights for their use in cancer research and clinical outcome prediction. Our findings suggest several priorities for future scFM development. Most evaluated models derive representations primarily from gene-expression information at the individual-cell level, whereas gene-regulatory relationships, pathways, tissue context, treatment exposure, cellular composition, and interactions among tumor-microenvironment compartments are represented inconsistently or left to downstream analysis. Similarly, most scFMs do not natively learn hierarchical representations connecting genes, cellular states, cell populations, and patients. Although the present analyses did not establish that these design characteristics caused the observed gap between cell-level embedding quality and patient-level prediction, they provide a plausible explanation for why strong biological organization of individual cells did not consistently translate into superior clinical performance. Increasing the number and diversity of patients represented in training data will remain essential, but greater dataset scale alone may not be sufficient. Future models should combine broader cancer datasets with structured biological knowledge, multimodal measurements, and multiscale learning objectives that explicitly connect molecular, cellular, microenvironmental, and patient-level information. Such models should be evaluated not only for cell-type recovery and batch mixing but also for their ability to preserve clinically relevant variation and produce interpretable predictions linked to biological pathways and cellular states.

Overall, our findings show that the value of an scFM is conditional on the intended task, the property of the representation being evaluated, and the method used to aggregate cell-level information. Foundation models captured biologically meaningful structure but did not consistently outperform strong conventional baselines; classification performance did not increase monotonically with model size; and cancer-specific or task-specific pretraining improved selected embedding properties without, by itself, establishing clinical utility. Under the evaluated protocols, end-to-end fine-tuning of Geneformer V1 and V2 did not consistently improve performance over frozen representations. Patient-level aggregation was also an integral component of the predictive pipeline: MIL most frequently achieved the highest AUPRC for conventional baselines and most scFMs, whereas the scGPT variants more often favored voting or embedding averaging. These results support an application-centered approach in which representation, adaptation, and aggregation strategies are evaluated jointly against the intended biological or clinical endpoint. To facilitate this process as the field evolves, scFM BenchScope combines automated model discovery with standardized, human-reviewed integration and reevaluation of additional models.

## Methods

### Single-cell foundation models and baseline approaches

We developed scFM BenchScope, a systematic framework for evaluating single-cell foundation models (scFMs) across cancer-relevant representation-learning and prediction tasks. We benchmarked 12 pretrained scFM variants together with three conventional baseline approaches across seven supervised tasks derived from publicly available single-cell RNA-sequencing datasets (Table 1 and Supplementary Table 1). The evaluated scFMs encompassed models with diverse architectures, pretraining objectives, and training corpora, with model sizes ranging from approximately 10 million to 600 million parameters and reported pretraining datasets spanning approximately 11 million to 167 million single-cell profiles.

The benchmark included four Geneformer variants: the first-generation Geneformer (GF-V1), the second-generation Geneformer (GF-V2), a deeper second-generation model (GF-V2-Deep), and a cancer-adapted second-generation model [GF-V2 (cancer)]. Geneformer V1 and V2 were additionally evaluated under supervised task-specific fine-tuning. Two scGPT checkpoints, a general-purpose model and a cancer-specific model [scGPT (cancer)], were included. The remaining scFMs - scFoundation, SCimilarity, scConcept, Nicheformer, STATE, and CellPLM - expanded the benchmark across alternative model architectures and pretraining strategies (Table 1).

Three non-foundation-model approaches were included as reference baselines. For the highly variable gene (HVG) baseline, the top 4,096 highly variable genes were used directly as expression features. For principal component analysis (PCA), expression values were scaled and projected onto 20, 50, or 100 principal components. For scVI, a variational autoencoder was trained on 4,096 HVGs using a negative-binomial observation model, and a 100-dimensional latent representation was extracted. Unless otherwise specified, fixed representations used for supervised classification were evaluated with a random forest classifier containing 100 trees. Exact preprocessing and model-specific parameters are provided in the Supplementary Methods and benchmark configuration files.

### Evaluation paradigms

We evaluated scFM representations under three complementary transfer settings: frozen inference, continual pretraining, and supervised fine-tuning. These settings were considered separately from the strategy used to aggregate individual cells into patient- or sample-level predictions.

In the frozen-inference setting, pretrained model weights were held fixed and cell-level representations were extracted without updating the foundation model on the evaluation dataset. For unsupervised analyses, these representations were evaluated directly for their ability to preserve annotated biological structure while limiting technical batch effects. For supervised analyses, the frozen embeddings were used as input features for task-specific classifiers. We refer to this setting as frozen-feature classification.

In continual-pretraining experiments, the pretrained encoder was further optimized on cells from the evaluation domain using the model’s self-supervised objective without using task labels. In supervised fine-tuning experiments, the pretrained encoder and task-specific prediction head were jointly optimized using labeled training data. Fine-tuning was performed independently within each cross-validation fold, and the corresponding held-out samples were not used for supervised parameter updates. Geneformer V1 and V2 were evaluated under this paradigm. Detailed optimization parameters are provided in the Supplementary Methods.

### Cancer prediction tasks

The supervised benchmark comprised seven cancer-relevant classification tasks spanning breast, lung, colorectal, and melanoma cohorts. These included: breast cancer subtype classification (ER-positive versus triple-negative breast cancer); lung adenocarcinoma treatment-status classification (treatment-naive versus tyrosine kinase inhibitor-treated); breast cancer treatment-status classification following anti-PD1 therapy; prediction of immune-checkpoint-blockade response in melanoma; classification of mismatch-repair-deficient versus mismatch-repair-proficient colorectal cancer; classification of exhausted versus non-exhausted T cells in breast cancer; and breast cancer treatment-status classification comparing treatment-naive and neoadjuvant chemotherapy samples. Dataset composition, cell populations, endpoint definitions, inclusion criteria, and individual cross-validation schemes are described in the Supplementary Methods and Supplementary Table 1.

### Evaluation of single-cell representations

We assessed whether pretrained representations preserved biologically meaningful cell-state structure while reducing unwanted batch-associated variation. A k-nearest-neighbor graph was constructed in embedding space using 15 neighbors and cosine distance. Agreement between unsupervised Leiden clusters and reference cell-type annotations was quantified using normalized mutual information (NMI) and adjusted Rand index (ARI), with Leiden resolution selected across a predefined search grid according to NMI. Biological conservation was further quantified using label average silhouette width (ASW), graph connectivity, k-nearest-neighbor label purity, and k-nearest-neighbor label prediction accuracy.

For datasets containing multiple batches, batch-associated structure was evaluated using batch ASW, principal-component regression (PCR), integration local inverse Simpson’s index (iLISI), cell-type local inverse Simpson’s index (cLISI), kBET, k-nearest-neighbor batch purity, and the predictability of batch identity from the representation. An aggregate biological-conservation score was computed from NMI, ARI, and label ASW.

To assess the stability of embedding-quality estimates, metrics were recomputed across 10 independently seeded stratified subsamples of up to 10,000 cells from each extracted embedding. Sampling preserved the joint distribution of batch and cell-type labels when both annotations were available; for datasets containing 10,000 cells or fewer, all available cells were used. Exact metric definitions and implementation details are provided in the Supplementary Methods.

### Patient- and sample-level prediction

Because clinical endpoints were defined at the patient or sample level whereas scFM representations were generated at the cell level, we compared three strategies for aggregating cellular information into a group-level prediction.

For mean pooling, cell embeddings belonging to the same patient or sample were averaged to produce a single representation, which was classified using a random forest model. For cell-level voting, a classifier was trained on individual cell embeddings using the group-level endpoint as the label. The final patient or sample class was determined by majority vote across cells, while the continuous prediction score used for AUC and AUPRC calculations was obtained by averaging cell-level predicted probabilities. Finally, for attention-based multiple-instance learning (MIL), all cells from a patient or sample were treated as a bag. A neural network learned cell-level transformations and attention weights and combined the resulting weighted cellular representations to generate a group-level prediction.

The downstream aggregation and classification strategies were evaluated using matched task-specific cross-validation partitions. All cells belonging to the same analysis unit used for a given task (patient, sample, or donor-timepoint) were kept together within the same fold.

### Classification performance and validation

Classification performance was assessed on held-out analysis units using area under the receiver-operating-characteristic curve (AUC), area under the precision-recall curve (AUPRC), accuracy, and macro-averaged F1 score, precision, and recall. Metrics were calculated independently for each cross-validation fold and summarized across folds.

Cross-validation used a task-specific grouping unit (patient, sample, or donor-timepoint). For tasks with substantial class imbalance, oversampling was restricted to the training partition. The number of folds and exact assignments were defined separately for each task and were held constant across model comparisons. Full split assignments are provided in Supplementary Table 9.

### Reproducibility

scFM BenchScope is available as open-source software at https://github.com/marakeby/scfm-benchscope. Benchmark experiments are specified through configuration files that record the dataset, model, preprocessing, task, transfer setting, classifier, and evaluation parameters. Resolved configurations and run-level results are retained with each experiment to support reproducibility and direct comparison across models. The software environments required by different pretrained models are isolated and dependency-pinned because individual scFMs rely on partially incompatible software stacks. Additional implementation and reproducibility details are provided in the Supplementary Methods

## Data availability

The processed datasets used for model evaluation are publicly available through the scFM [BenchScope Evaluation Data repository](https://www.dropbox.com/scl/fo/8pnyn6izq475vobdsw8ns/AGd6mUqU36JzrVne4mk0XMY?rlkey=48rx79ol75rzgxm20dmohf98i&st=h8vk5fwp&dl=0). The raw datasets are available from the repositories associated with their original publications. Dataset accession information and corresponding references are provided in the Supplementary Methods.

## Code availability

The source code, configuration files, and analysis workflows used to perform the benchmark are publicly available in the scFM BenchScope GitHub repository. Benchmark results and interactive visualizations are available through the scFM BenchScope dashboard

## Acknowledgements

This work was supported by:

NIH: SPORE P50CA272390, P01CA228696, R01CA278980 (E.M.V.)

DOD: W81XWH-21-PCRP-DSA, and DOD HT94252410415 (E.M.V.)

DOD CDMRP award (HT9425-23-1-0023) (HAE)

Damon Runyon Quantitative Biology Fellowship (A.R.)

McDonough Fellowship (A.R.)

Mark Foundation Emerging Leader Award (E.M.V.)

Quad Fellowship (S.J.)

## Author Disclosures

### Competing interests

EMV: Advisory/Consulting: Enara Bio, Manifold Bio, Monte Rosa, Novartis Institute for Biomedical Research, Serinus Bio, TracerBio Research support: Novartis, BMS, Sanofi, NextPoint

Equity: Tango Therapeutics, Genome Medical, Genomic Life, Enara Bio, Manifold Bio, Microsoft, Monte Rosa, Riva Therapeutics, Serinus Bio, Syapse, TracerDx

Travel reimbursement: None

Speaking Fees: TD Cowen

Patents: Institutional patents filed on chromatin mutations and immunotherapy response, and methods for clinical interpretation; intermittent legal consulting on patents for Foaley & Hoag

Editorial Boards: *Science Advances*

HE, AR, and SJ declare no competing interests

## Author Contributions

Conceptualization: HAE, AR, and EMV;

Methodology: HAE, AR, and SJ;

Software: HAE, and AR;

Investigation: HAE, AR, SJ, and EMV;

Writing, original draft: HAE, AR, and EMV;

Writing, review and editing: HAE, AR, SJ, and EMV;

Visualization: HAE, and AR;

Supervision: HAE and EMV;

Funding acquisition: HAE, AR, and EMV

## Supplementary Information

### Empirical Evaluation of Single-Cell Foundation Models for Predicting Cancer Outcomes

#### Data

We analyzed six publicly available single-cell RNA-sequencing datasets spanning breast cancer, lung adenocarcinoma, colorectal cancer, and melanoma. Dataset-specific inclusion criteria, annotations, and supervised endpoints are described below.

#### Bassez et al. 2021^1^ - breast cancer

Raw data (.rds objects) and accompanying metadata (.csv files) were obtained from the Lambrechts laboratory single-cell resource. The study included ER-positive, HER2-positive, and triple-negative breast cancer (TNBC) samples collected from treatment-naive patients and from patients treated with anti-PD1 therapy or chemotherapy plus anti-PD1 therapy. HER2-positive patients were excluded because of the small number of available cases.

The dataset contributed four supervised analyses:

i. treatment-naive ER-positive versus treatment-naive TNBC;
ii. treatment-naive versus anti-PD1-treated samples;
iii. exhausted versus non-exhausted T cells; and
iv. treatment-naive versus neoadjuvant-chemotherapy-treated samples.

Author-provided annotations were used to identify tumor cells, T cells, B cells, and other cell populations. The dataset was also used for unsupervised evaluation of cell-type organization in the learned embeddings.

#### Kim et al. 2020 ^2^ - lung adenocarcinoma

Raw UMI counts (GSE131907_Lung_Cancer_raw_UMI_matrix.txt.gz) and cell-type annotations (GSE131907_Lung_Cancer_cell_annotation.txt.gz) were obtained from GEO accession GSE131907. Patient-level clinical metadata were obtained from Supplementary Table S1 of the original publication. Only samples obtained from primary tumor sites were retained; pleural-effusion and metastatic-site samples were excluded. Author-provided tumor-cell annotations were used.

#### Maynard et al. 2020 ^3^- lung adenocarcinoma

Processed single-cell objects were obtained from the public data repository associated with the original study. NI01_Nonimmune_Seurat_object_annotated.RData contained epithelial cells, and NI05_all_epithelial_annotated_normal_and_tumor.RData provided tumor-versus-normal annotations. Additional clinical metadata were obtained from Supplementary Table S1 of the publication. Only primary-tumor samples from patients diagnosed with lung adenocarcinoma were retained for analysis.

#### Qian et al. 2020 ^4^- lung adenocarcinoma

Raw lung adenocarcinoma data were obtained from the Lambrechts laboratory Pan-Cancer Blueprint of the Tumour Microenvironment resource. Author-provided tumor-cell annotations were used, and additional tumor-stage information was obtained from Supplementary Table S1 of the original publication.

#### Meta-analysis of lung adenocarcinoma datasets

The Kim, Maynard, and Qian lung adenocarcinoma datasets were analyzed jointly for treatment state classification (treatment-naive versus tyrosine kinase inhibitor (TKI)-treated disease)

#### Pelka et al. 2021^5^ - colorectal cancer

The raw expression count matrix was obtained from GEO accession GSE178341, and corresponding cell-level metadata and annotations were obtained from the Broad Institute Single Cell Portal (SCP1162). The dataset contains 371,223 cells collected from primary, treatment-naive colorectal tumors and adjacent normal colon tissue from 62 patients, including 34 mismatch-repair-deficient (MMRd) and 28 mismatch-repair-proficient (MMRp) patients. MMR status, tissue identity, patient identity, and cell-type annotations were obtained from the accompanying metadata and Supplementary Table S1A of the original publication. Tumor and adjacent-normal samples from patients with primary colorectal cancer were retained for downstream analysis.

#### Sade-Feldman et al. 2018 ^6^- melanoma

The processed expression matrix and associated cell-level metadata were obtained from GEO accession GSE120575. The study profiled 16,291 CD45-positive immune cells from 48 metastatic melanoma biopsies and included patients treated with immune checkpoint blockade. Biopsies were annotated by clinical response and treatment timing (pre-treatment or on-treatment). Using the response definitions reported by the original study, responders had complete or partial responses and non-responders had stable or progressive disease. Only pre-treatment samples were used for evaluation of baseline predictors of therapeutic response.

### Benchmark implementation

scFM BenchScope was developed as a modular evaluation framework for comparing single -cell foundation models (scFMs) and conventional representation-learning approaches under a common experimental protocol. Individual experiments are defined by configuration files specifying the dataset, model, preprocessing procedure, downstream task, training paradigm, classification strategy, and evaluation parameters. Reusable configuration components are combined to generate a fully resolved experiment specification before execution.

The evaluation workflow consists of data loading and cohort selection, quality control and preprocessing, model-specific input preparation, embedding extraction or model fine-tuning, representation-quality assessment when applicable, supervised classification, and generation of standardized run-level outputs. Dataset loaders, feature extractors, foundation-model adapters, and classifiers implement common interfaces, allowing additional models to be incorporated without modification of downstream evaluation code.

### Software environment and reproducibility

The framework is distributed as the open-source Python package scfm-cancer-eval and requires Python 3.10 or later. The package provides the scfm-eval command-line interface for experiment execution, configuration validation, and report generation. Core dependencies include NumPy, pandas, AnnData, Scanpy, scikit-learn, PyTorch, scIB, and related scientific Python packages.

Because pretrained scFMs frequently require incompatible versions of deep-learning and model-specific dependencies, model families were executed in isolated Pixi environments with dependency versions recorded by lockfiles. A default environment was used for common preprocessing, conventional baselines, downstream classifiers, and evaluation. Containerized execution can reproduce the same environments while keeping datasets and pretrained weights external to the software image.

Each benchmark run stores the resolved experiment configuration together with the resulting metrics and associated artifacts. Random seeds are explicitly specified in experiment configurations and model adapters and are retained with the run metadata. Where deterministic execution is supported by the corresponding libraries and hardware, deterministic settings are enabled. Cross-validation assignments are stored separately and reused across model comparisons, ensuring that models evaluated on the same task use identical train-test partitions.

Run-level outputs contain performance metrics and provenance information sufficient to associate a result with the corresponding software revision and configuration.

### Dataset preprocessing and cohort definition

Single-cell datasets were represented as AnnData objects containing expression measurements together with task-specific cell, patient, sample, treatment, and batch metadata. Before model-specific processing, each analysis applied the inclusion and exclusion criteria defined for the corresponding task. These filters restricted analyses to the relevant cancer type, anatomical site, treatment setting, and cellular compartment.

For analyses using the standard preprocessing workflow, cells with fewer than 10 detected genes and genes detected in fewer than 10 cells were excluded. Library-size normalization to 10,000 counts per cell followed by log1p transformation was used for methods requiring normalized expression input. When highly variable genes were required, 4,096 genes were selected using the Seurat highly-variable-gene procedure, with batch-aware selection enabled when specified by the experiment.

These preprocessing operations were not imposed uniformly on models whose published inference procedures require alternative input representations. After common cohort filtering, each model adapter applied the transformations required by the corresponding pretrained model, including procedures such as gene ranking, tokenization, expression binning, or model-specific gene-vocabulary mapping. Thus, models were evaluated on matched biological cohorts while retaining the native input conventions of each pretrained model.

### Baseline representations

Three conventional approaches were used as reference baselines. For the HVG baseline, expression values for the 4,096 selected highly variable genes constituted the representation directly. For PCA, expression values were scaled before dimensionality reduction, with scaled values clipped to a maximum absolute value of 10; PCA representations containing 20, 50, or 100 components were evaluated. For scVI, a variational autoencoder was trained on the evaluation dataset using the selected highly variable genes and a negative-binomial gene-expression likelihood. The benchmark configuration used a two-layer encoder and a 100-dimensional latent representation.

For the supervised benchmark, these baseline representations were fitted on the evaluation cohort before downstream cross-validation of the patient- or sample-level classifier. Clinical endpoint labels were not used to fit HVG selection, PCA, or scVI; however, unlabeled expression profiles from subsequently held-out units could contribute to the cohort-level representation fit. Accordingly, these baseline analyses represent transductive unsupervised representation learning followed by cross-validated supervised classification. Exact model and preprocessing parameters are retained in the resolved configurations accompanying the benchmark results.

### Foundation-model representation extraction

Cell-level representations were generated from published pretrained checkpoints for Geneformer, scGPT, scFoundation, CellPLM, SCimilarity, Nicheformer, scConcept, and STATE. Each model-specific adapter converted the filtered single-cell dataset to the representation required by the corresponding model and returned one fixed-length embedding per cell. These cell-by-feature matrices formed the input to unsupervised representation-quality analyses and, in frozen-feature experiments, to downstream classifiers. The extent to which pretrained model parameters were updated on the evaluation dataset depended on the transfer paradigm: frozen inference, continual pretraining, or supervised fine-tuning.

#### Frozen inference

In frozen-inference experiments, all pretrained model parameters were held fixed. Cells were processed using the published model checkpoint and corresponding model-specific input transformation, and the resulting representations were extracted without using labels from the benchmark task.

For unsupervised representation analyses, these embeddings were evaluated directly for biological conservation and batch-associated structure. For supervised prediction tasks, the embeddings were treated as fixed features and a task-specific classifier or aggregation model was trained using labeled data from the corresponding training partition.

#### Continual pretraining

Continual pretraining was used to evaluate whether unsupervised adaptation to the target single - cell domain improved the biological organization of pretrained representations. For Geneformer experiments, the pretrained model was further optimized using its self-supervised pretraining objective on cells from the target training dataset without using clinical or biological endpoint labels. The model encoder was then used to extract representations for subsequent evaluation using the same metrics applied to the corresponding frozen model. No supervised classification head was optimized during continual-pretraining experiments. For Geneformer, continual pretraining used a linear learning-rate schedule, a learning rate of 10^−4^, 6 epochs, batch size 16, weight decay of approximately 5 ∗ 10^−4^ to 10^−3^, and 1,000 warmup steps. Exact parameters for individual runs are retained in the resolved experiment configurations.

#### Supervised Geneformer fine-tuning

Geneformer V1 and V2 were additionally evaluated using supervised end-to-end fine-tuning. Fine-tuning was performed independently within each cross-validation fold. Only cells belonging to training analysis units were available for parameter optimization, and the pretrained encoder and patient-level prediction head were jointly updated.

Each patient was represented as a collection of cells and training used multiple-instance learning to obtain a patient-level prediction from cell-level transformer representations. The implementation supports sampling up to 1,000 cells per patient and processing cells in bags of up to 500. All Geneformer encoder layers were eligible for optimization (freeze_layers = 0). Optimization used AdamW with a learning rate of 2 ∗ 10^−5^ and weight decay of 0.01, with gradient checkpointing enabled to reduce memory requirements. Geneformer V2 (104M) was trained for 3 epochs and Geneformer V1 (10M) for 5 epochs. Model-specific sequence-length settings were defined in the resolved configuration for each checkpoint. Fine-tuned models were evaluated only on the held-out fold after training.

### Evaluation of embedding quality

Representation quality was evaluated from cell-level embeddings using metrics designed to quantify preservation of biological structure and removal of technical batch-associated structure. A k-nearest-neighbor graph was constructed from each embedding using 15 neighbors and cosine distance. Leiden clustering was performed across a predefined resolution grid, and the resolution maximizing normalized mutual information (NMI) with the annotated biological labels was selected for calculation of cluster-based metrics.

Biological conservation was quantified using normalized mutual information (NMI), adjusted Rand index (ARI), label average silhouette width (ASW), graph connectivity, k-nearest-neighbor label purity, and k-nearest-neighbor label prediction accuracy. An aggregate biological score was calculated as the mean of NMI, ARI, and label ASW.

For datasets containing more than one batch, batch-associated structure was additionally evaluated using batch ASW, principal-component regression (PCR) with respect to batch, integration local inverse Simpson’s index (iLISI), cell-type local inverse Simpson’s index (cLISI), kBET, k-nearest-neighbor batch purity, and batch-prediction accuracy. Together, these measures characterize complementary aspects of local cell-type conservation, neighborhood mixing, and the amount of batch information retained in the representation.

To quantify the stability of embedding metrics, the post-embedding analysis was repeated across 10 independently seeded subsampling runs. Each run selected up to 10,000 cells from the already extracted embedding using a different random seed. Sampling was stratified on the joint batch and cell-type label when both were available, or on batch when only batch information was available. Sampling was performed without replacement; therefore, this procedure is more precisely described as repeated stratified subsampling rather than classical bootstrap resampling. For datasets containing 10,000 cells or fewer, the complete set of cells was evaluated in each run. Metrics were recomputed independently for each subsample. Two-dimensional projections were generated from the extracted representations for qualitative visualization but were not used to calculate classification performance.

### Patient- and sample-level classification

Clinical and molecular endpoints were defined at the patient, sample, or donor-timepoint level, whereas foundation-model embeddings were generated for individual cells. We therefore evaluated three aggregation approaches for converting cell-level representations into predictions at the endpoint level.

**1. Embedding Averaging.** Cell embeddings were averaged within each analysis unit to obtain a single feature vector. A random forest classifier was trained on these aggregate representations.
**2. Cell-level voting.** A random forest classifier was trained using individual cell embeddings, with each cell assigned the endpoint of its corresponding analysis unit. At inference, cell-level predicted classes were combined by majority vote to obtain the final patient- or sample-level class. For binary endpoints, the continuous group-level score used for AUC and AUPRC calculation was obtained by averaging cell-level predicted probabilities for the positive class.
**3. Attention-based multiple-instance learning (MIL).** Each patient or sample was represented as a bag of cell embeddings. The MIL network first transformed individual cell representations through a neural feature encoder and then learned temperature-scaled attention weights for the cells in each bag. A weighted sum of the transformed cell representations generated the bag representation, which was passed to a classification network to produce the endpoint-level prediction. The benchmark configuration used hidden and attention dimensions of 128 and 64, respectively, dropout of 0.3, Adam optimization with a learning rate of 10^−4^ and weight decay of 10^−5^, and training for up to 500 epochs with early stopping. Attention weights were retained to allow ranking of cells according to their contribution to the bag representation.

### Classification metrics

Classification performance was evaluated using predictions for held-out analysis units. For binary outcomes, performance was summarized using the area under the receiver-operating-characteristic curve (AUC) and area under the precision-recall curve (AUPRC), together with accuracy, macro-averaged F1 score, precision, and recall. Metrics were calculated separately within each cross-validation fold and then summarized across folds. Sample and cell-level predictions, fold-level performance metrics, and corresponding experimental configurations were retained for each run.

### Data splits and validation

Cross-validation was performed using the task-specific independent unit defined for each analysis. All cells belonging to the same unit/sample were assigned to the same fold. The same predefined fold assignments were reused across models and aggregation strategies to permit paired model comparisons. Test sets were never oversampled. For tasks with substantial class imbalance, oversampling was applied only to the training partition. Exact assignments are provided in Supplementary Table 9.

**Task 1: LUAD (treatment-naive / TKI).** Four-fold cross-validation was performed on 8 samples (sample_id); each fold contained 6 training samples and 2 test samples. Every test set included one treatment-naive and one TKI-treated sample.

**Task 2: BRCA (ER-positive / TNBC subtypes).** Cancer cells from pretreatment samples were analyzed using five-fold cross-validation on 28 patients. Each fold included 22-23 training patients and 5-6 test patients. Test sets included both ER-positive and TNBC patients.

**Task 3: BRCA (treatment-naive / anti-PD1).** T cells were labeled by pre-versus post-treatment status. Splits were defined on donor-timepoint units (donor_id_pre_post; 62 units from 31 donors, each contributing one Pre and one Post sample). Five-fold cross-validation was used; each fold contained 49-50 units in training and 12-13 units in testing, with both pre- and post-treatment samples represented in the test fold.

**Task 4: Melanoma (immune-checkpoint-blockade response).** Pretreatment lesions were labeled as responder (R) versus non-responder (NR). Five-fold cross-validation was performed on 19 samples, with 15-16 samples in training and 3-4 in testing per fold. Every test fold contained both responders and non-responders.

**Task 5: CRC (MMRd versus MMRp).** Tumor-infiltrating T cells were evaluated using five-fold cross-validation on 58 patients. Each fold included 46-47 patients in training and 11-12 in testing, with both MMRd and MMRp patients represented in the test fold.

**Task 6: BRCA (exhaustion status: NE / E).** T cells from pretreatment samples were evaluated using five-fold cross-validation on 29 patients. Each fold included 23-24 unique patients in training and 5-6 in testing. Test sets contained both non-exhausted and exhausted samples. Training folds were oversampled to mitigate class imbalance; test sets were not oversampled.

**Task 7: BRCA (treatment-naive / chemotherapy).** Cancer cells were evaluated using five-fold cross-validation on 39 patients. Each fold included 31-32 unique patients in training and 7-8 in testing. Test sets included both treatment-naive and neoadjuvant-chemotherapy patients. Training folds were oversampled to mitigate class imbalance; test sets were not oversampled.

### Evaluated models

We compared eight pretrained single-cell foundation models, using the authors’ public checkpoints, against three classes of classical baselines (PCA, highly variable genes, and scVI). Where a family released more than one checkpoint (Geneformer, scGPT), we evaluated each variant under the same protocol. Pretrained models were used as frozen encoders unless otherwise noted (Geneformer continual pretraining and supervised fine-tuning; see below). Table 1 in the main paper summarizes the evaluated models.

#### Geneformer ^7^

Geneformer is a BERT-style transformer that represents each cell as a rank-ordered sequence of genes (rank-value encoding) and is pretrained with masked language modeling: 15% of gene tokens are masked and the model predicts gene identity from the remaining context. We evaluated four published checkpoints from the same series. Geneformer V1 (10 million parameters; 6 layers, embedding dimension 256, 4 attention heads, context 2,048) was pretrained on Genecorpus-30M (∼30 million human single-cell transcriptomes, excluding high-mutational-burden malignant cells). Geneformer V2-104M (104 million parameters; 12 layers, dimension 768, 12 heads, context 4,096) and V2-316M (316 million parameters; 18 layers, dimension 1,152, 18 heads, context 4,096) were pretrained on Genecorpus-104M (∼104 million non-cancer human transcriptomes). A cancer-domain variant (V2-104M CLcancer) continues V2-104M pretraining on ∼14 million cancer transcriptomes.

#### scGPT ^8^

scGPT is a generative pretrained transformer that tokenizes gene identity together with binned expression values and uses a specialized attention mask so that masked gene expression can be predicted from unmasked genes while a cell embedding is learned jointly. The published backbone has 12 transformer blocks, 8 attention heads, and embedding dimension 512 (approximately 53 two-step masked gene-expression objective (gene-prompt generation plus cell-prompt generation; mean-squared error on masked expression bins). We evaluated the whole-human checkpoint, pretrained on 33 million normal (non-disease) human cells from CELLxGENE (51 organs/tissues, 441 studies), and the pan-cancer checkpoint, pretrained on 5.7 million cells spanning multiple cancer types. Inference used a maximum sequence length of 2,048 tokens and 51 expression bins.

#### scFoundation ^9^

scFoundation is a 100-million-parameter model built on the xTrimoGene asymmetric encoder–decoder: the encoder attends only to non-zero, non-masked gene tokens, while the decoder reconstructs the full 19,264-gene profile. It was pretrained on more than 50 million human scRNA-seq profiles with a read-depth-aware masked reconstruction task (30% of genes masked, including zeros): the model predicts masked expression given source and target total-count indicators, using mean-squared error on masked positions. We used the published 100-million-parameter checkpoint (01B-resolution) and pooled encoder outputs as cell embeddings.

#### CellPLM ^10^

CellPLM treats cells as tokens and tissues as sentences, so that cell–cell context is modeled explicitly. A gene-expression embedder feeds a linear-complexity Flowformer encoder; a Gaussian-mixture variational latent space regularizes noisy transcriptomes, and a decoder reconstructs masked gene expression. Pretraining optimizes a denoising evidence lower bound (mean-squared reconstruction of masked genes plus KL divergence to the mixture prior). The released 85-million-parameter checkpoint (20231027_85M; 82.4 million parameters in the paper) was pretrained on more than 9 million scRNA-seq cells and about 2 million spatially resolved cells (∼11 million cells in total).

#### Nicheformer ^11^

Nicheformer is a transformer encoder pretrained jointly on dissociated and spatially resolved transcriptomics. The model has 12 encoder layers, 16 attention heads, embedding dimension 512, feed-forward size 1,024, and a 1,500-token context (49.3 million parameters). Gene-rank tokens are concatenated with modality, species, and assay context tokens, including a niche token for spatial samples. Pretraining uses BERT-style masked language modeling (15% of tokens masked). The training corpus is SpatialCorpus-110M: more than 110 million cells (∼57 million dissociated and ∼54 million spatial) from human and mouse across 73 tissues. We used the public Hugging Face checkpoint (theislab/Nicheformer) in dissociated, human mode.

#### SCimilarity ^12^

SCimilarity learns a metric embedding for cross-study cell search rather than a masked language model. A fully connected encoder maps 28,231 genes through hidden layers of width 1,024–1,024–1,024 to a 128-dimensional, unit-normalized embedding (approximately 31 million encoder parameters); a decoder is used only during training. The training objective combines a supervised triplet loss (same Cell Ontology type, different studies, versus a different type) with mean-squared reconstruction of the input profile (relative triplet weight (b = 0.001), margin 0.05). The encoder was trained on 7.9 million annotated cells from 56 studies; the associated search atlas comprises 23.4 million profiles from 412 studies. We used SCimilarity v1.1.

#### scConcept ^13^

scConcept replaces gene-level reconstruction with contrastive pretraining of cell embeddings: two augmented views of the same cell are pulled together and contrasted with other cells, yielding representations that are intended to be invariant to count distribution and gene-panel choice. The architecture is a transformer encoder; the loss is a cell-level contrastive (identification) objective rather than masked language modeling. We used the Corpus-30M checkpoint, pretrained on more than 30 million human scRNA-seq profiles (approximately 30 million parameters; 8 transformer layers, hidden size 512, 8 attention heads). Cell embeddings were the mean of gene-token representations.

#### STATE ^14^

STATE comprises a State Embedding (SE) model of cellular identity and a State Transition model of perturbation responses. We evaluated only SE: a bidirectional transformer encoder (16 layers, 16 attention heads, hidden size 2,048) with an MLP decoder, totaling about 600 million parameters (checkpoint SE-600M, epoch 16). SE was trained on 167 million unperturbed human cells (standardized to 19,790 protein-coding genes). Training uses masked gene reconstruction (mask rate 0.2) with an energy-based (tabular) loss that matches predicted and observed expression distributions, rather than a standard token-level cross-entropy.

#### Classical baselines

PCA, HVG features, and scVI were fit on each evaluation cohort and are not pretrained foundation models. PCA used Scanpy after scaling with 20, 50, or 100 components. HVG used the top 4,096 Seurat-flavor genes as the representation. scVI ^15^ is a hierarchical Bayesian variational autoencoder with a negative-binomial observation model; we trained a two-layer encoder with a 100-dimensional latent space for 10 epochs on the evaluation data, optimizing the evidence lower bound (negative-binomial reconstruction plus KL regularization of the approximate posterior). Parameter counts for scVI depend on the number of genes in each dataset.

### Model discovery and extensibility

The benchmark includes an assisted model-discovery workflow that surveys public literature and software repositories for newly released scFMs and records candidate models together with their publications, source repositories, pretrained weights, and available metadata. Integration into the benchmark requires a model-specific adapter, a reproducible dependency specification, pinned model resources, benchmark configuration, successful execution, and scientific review of the resulting run. An integration agent is used to automate the evaluation process with input from human reviewer to accept/decline/modify the proposed plan and implementation. Automated discovery and integration agents therefore support benchmark maintenance while remaining separate from the core evaluation framework.

### Results reporting and interactive resource

Standardized run outputs are aggregated to generate the scFM BenchScope reporting resource and companion interactive dashboard (https://marakeby.github.io/scfm-benchscope/). The dashboard summarizes embedding-quality and classification results and provides a searchable catalog of evaluated scFMs together with a metric dictionary. The reporting layer consumes the same structured run outputs used for manuscript analyses, reducing the need for manual transcription of benchmark results.

### Code availability

Source code for scFM BenchScope, benchmark configurations, model adapters, data-loading procedures, evaluation routines, and documentation is publicly available at https://github.com/marakeby/scfm-benchscope. The repository contains instructions for reproducing model-specific software environments and executing benchmark experiments.

---

**Supplementary Figure 1.**
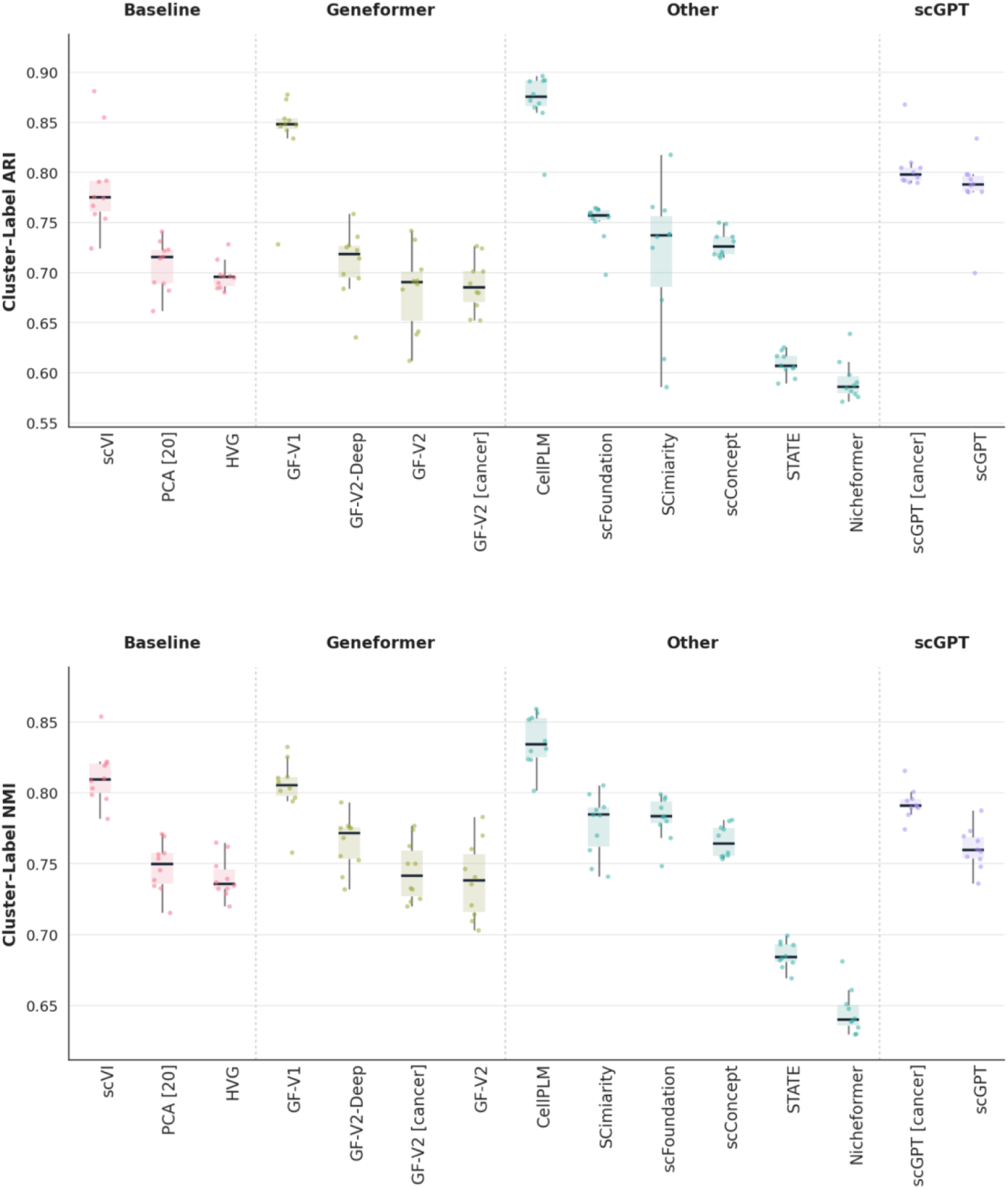
Embedding quality evaluation. Cluster–label ARI (top) and NMI (bottom) were calculated between Leiden clusters and annotated cell types. For each embedding, 10 subsamples of 10,000 cells were drawn without replacement and stratified by batch and cell type. Leiden clustering was performed on a 15-nearest-neighbor graph using cosine distance, with the resolution selected to maximize NMI. Higher values indicate stronger cluster–label agreement. Fifteen embedding configurations are grouped as Baseline, Geneformer, Other, and scGPT and ordered within each group by median score. Boxes show the interquartile range, central lines show medians, and points represent individual subsampling runs

**Supplementary Figure 2.**
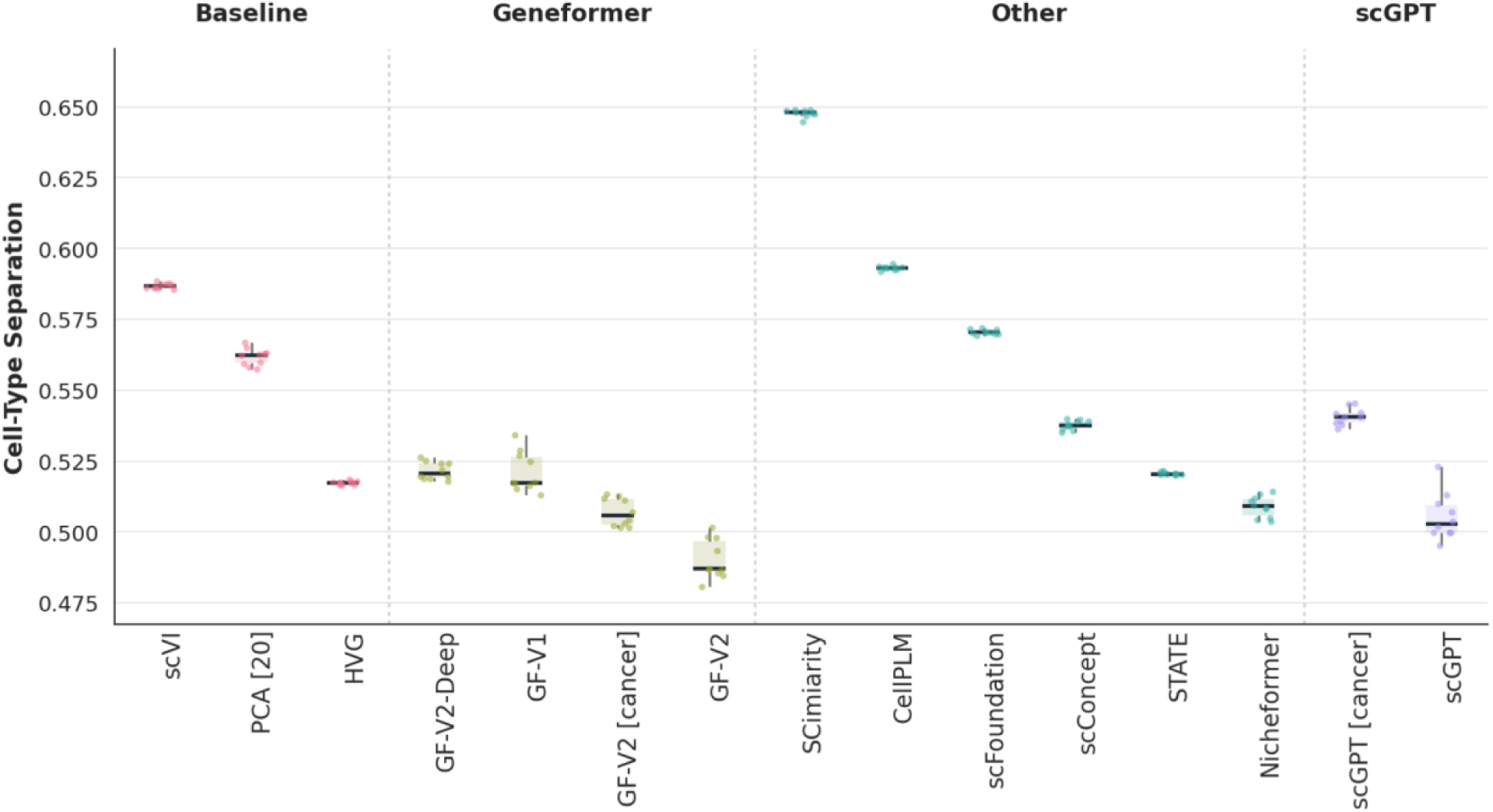
Cell-type separation. Cell-type separation was measured using the average silhouette width (ASW) of annotated cell types in embedding space, based on Euclidean distances. Scores were calculated across 10 stratified subsamples of 10,000 cells drawn without replacement. Higher values indicate more compact and distinct cell-type groups. Fifteen embedding configurations are grouped as Baseline, Geneformer, Other, and scGPT and ordered within each group by median score. Boxes show the interquartile range, central lines show medians, and points represent individual subsampling runs.

**Supplementary Figure 3.**
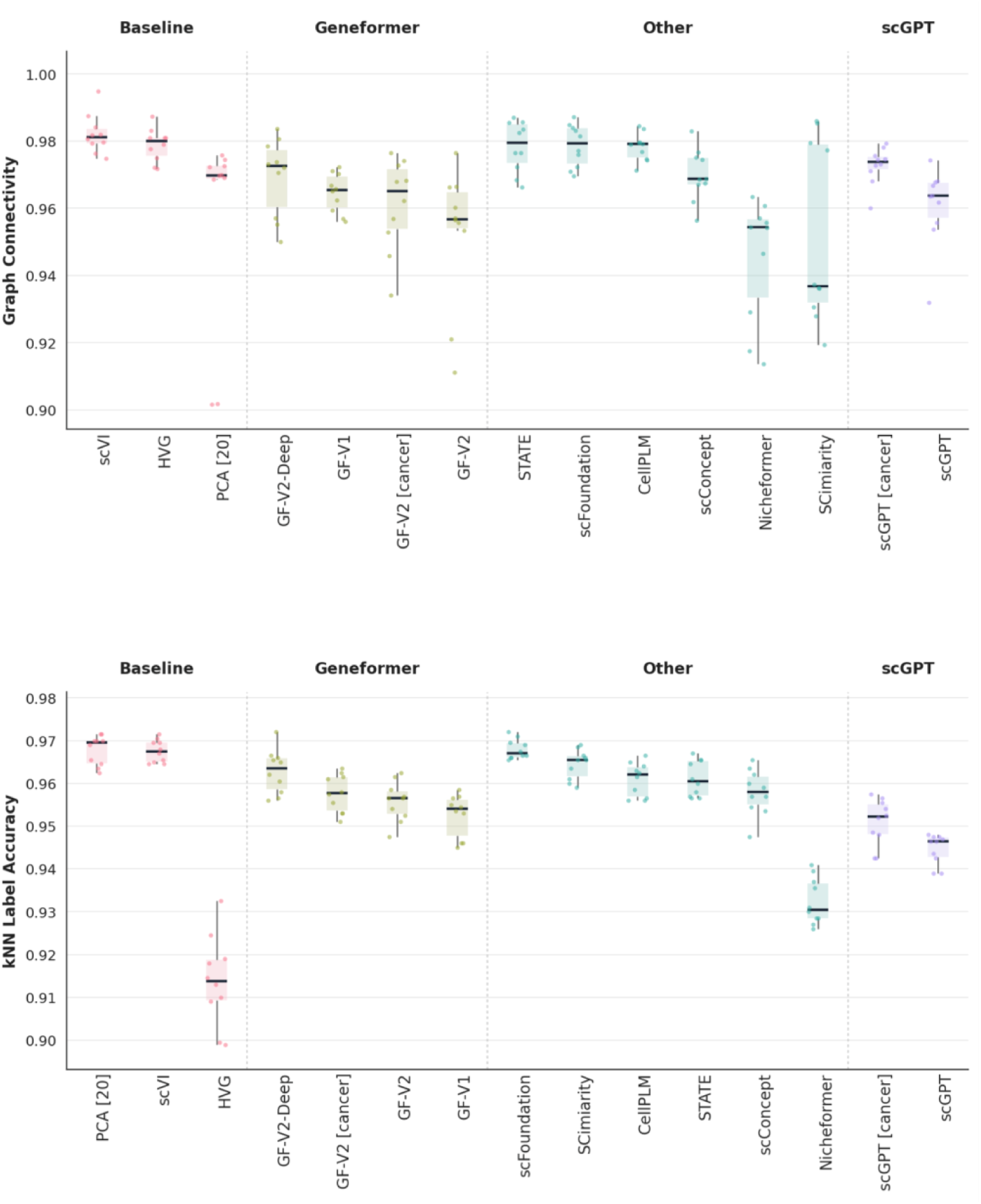
Graph connectivity and kNN label accuracy. **Top:** Graph connectivity measures the proportion of cells within each annotated cell type belonging to its largest connected component in the embedding-derived neighbor graph, averaged across cell types. **Bottom:** kNN label accuracy measures cell-type prediction accuracy using a distance-weighted 15-nearest-neighbor classifier trained on 80% of cells and evaluated on a stratified 20% holdout set. Scores were calculated across 10 stratified subsamples of 10,000 cells drawn without replacement. Higher values indicate better preservation of cell-type structure. Fifteen embedding configurations are grouped by model family and ordered within each group by median score. Boxes show the interquartile range, central lines show medians, and points represent individual subsampling runs.

**Supplementary Figure 4.**
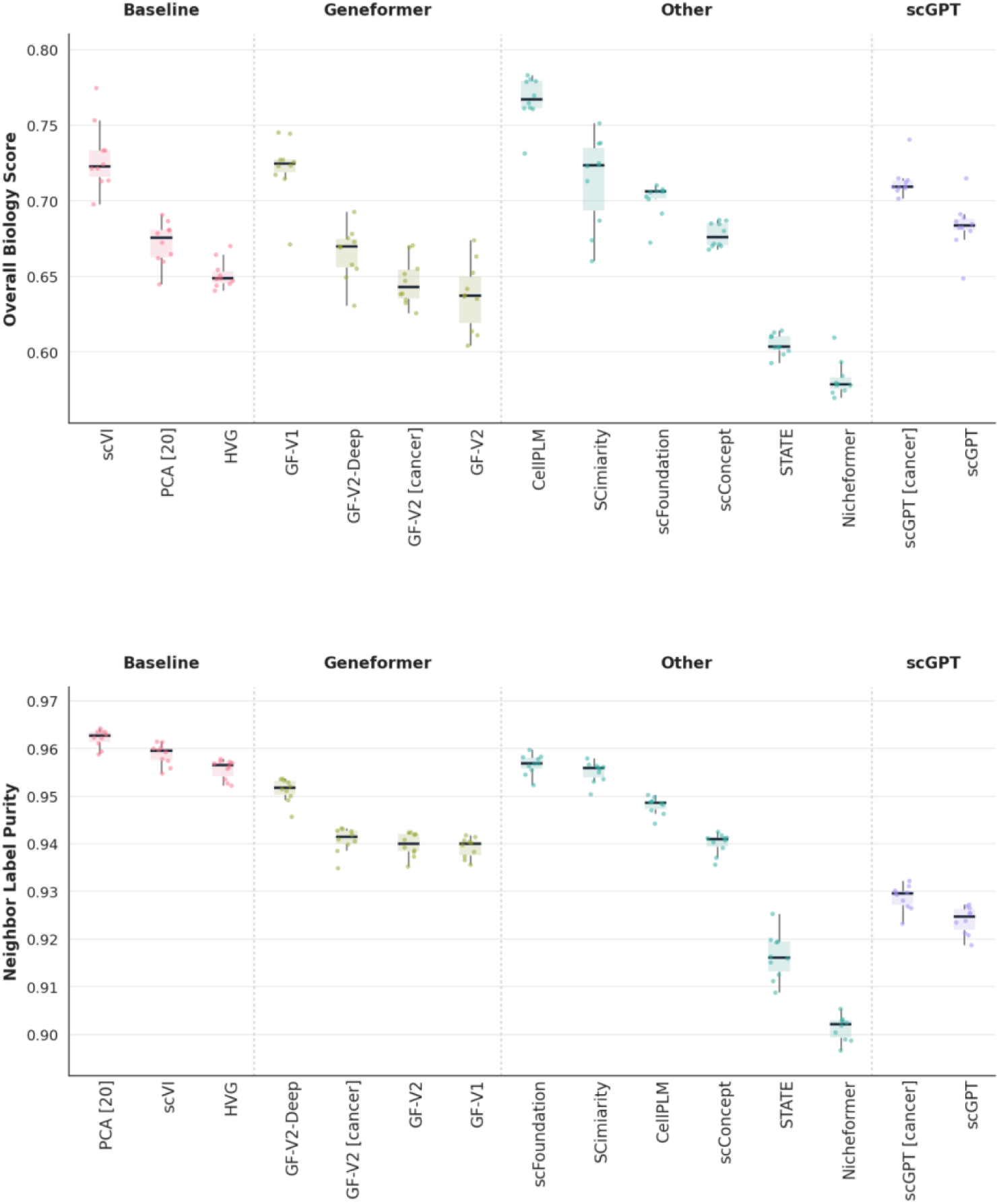
Overall biological conservation and neighbor-label purity. **Top:** The overall biology score is the mean of cluster–label NMI, cluster–label ARI, and cell-type silhouette width, summarizing the preservation of cell-type structure. **Bottom:** Neighbor-label purity is the mean proportion of each cell’s 15 nearest neighbors that share its annotated cell-type label. Scores were calculated across 10 stratified subsamples of 10,000 cells drawn without replacement. Higher values indicate better preservation of biological structure. Fifteen embedding configurations are grouped by model family and ordered within each group by median score. Boxes show the interquartile range, central lines show medians, and points represent individual subsampling runs.

**Supplementary Figure 5.**
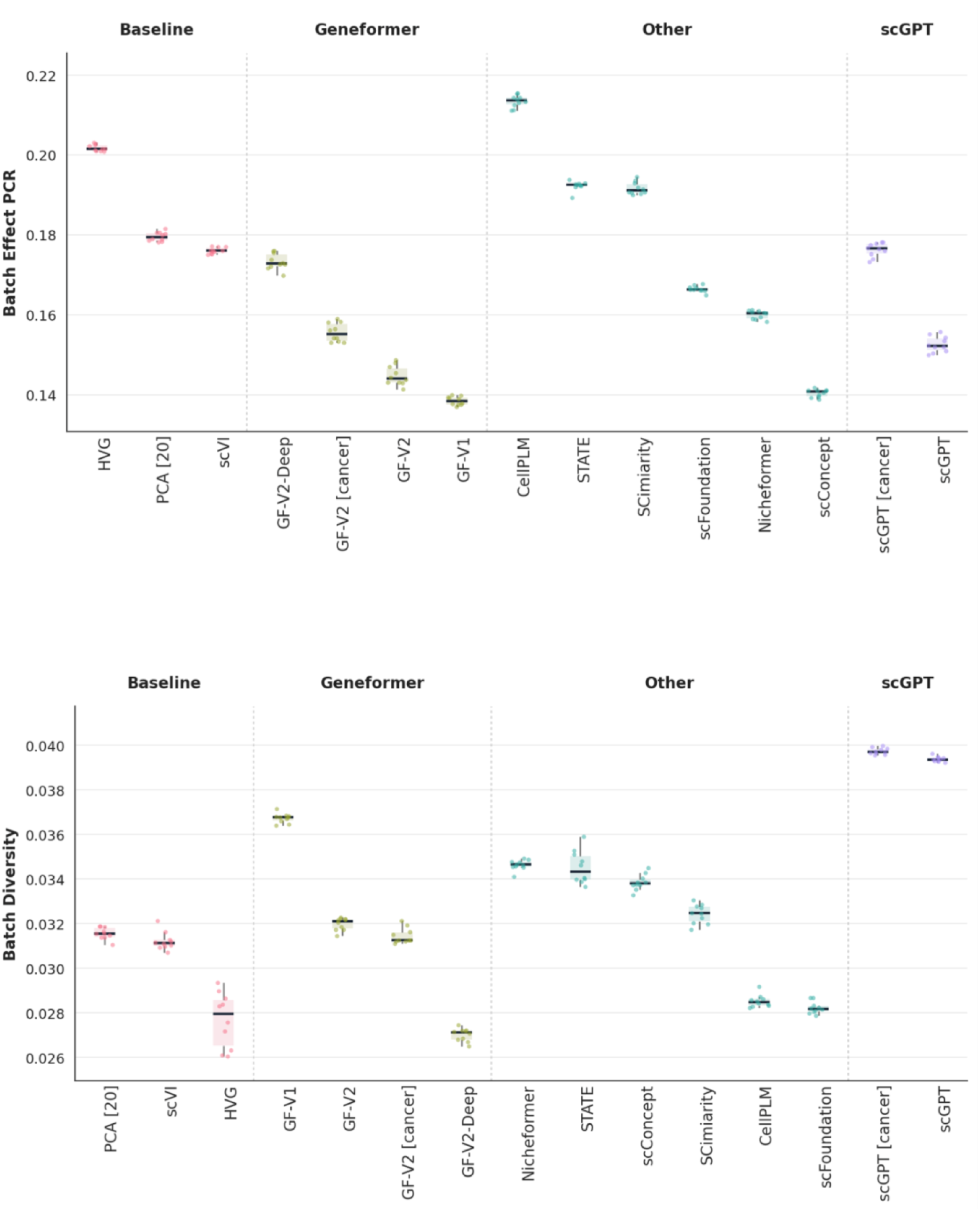
Batch-effect PCR and batch diversity. **Top:** Batch-effect principal component regression (PCR) measures the proportion of variation in the embedding associated with batch identity; lower values indicate weaker batch effects. **Bottom:** Integration Local Inverse Simpson’s Index (iLISI) measures batch diversity within local cell neighborhoods; higher values indicate greater batch mixing. Scores were calculated across 10 stratified subsamples of 10,000 cells drawn without replacement. Fifteen embedding configurations are grouped by model family and ordered within each group by median score. Boxes show the interquartile range, central lines show medians, and points represent individual subsampling runs.

**Supplementary Figure 6.**
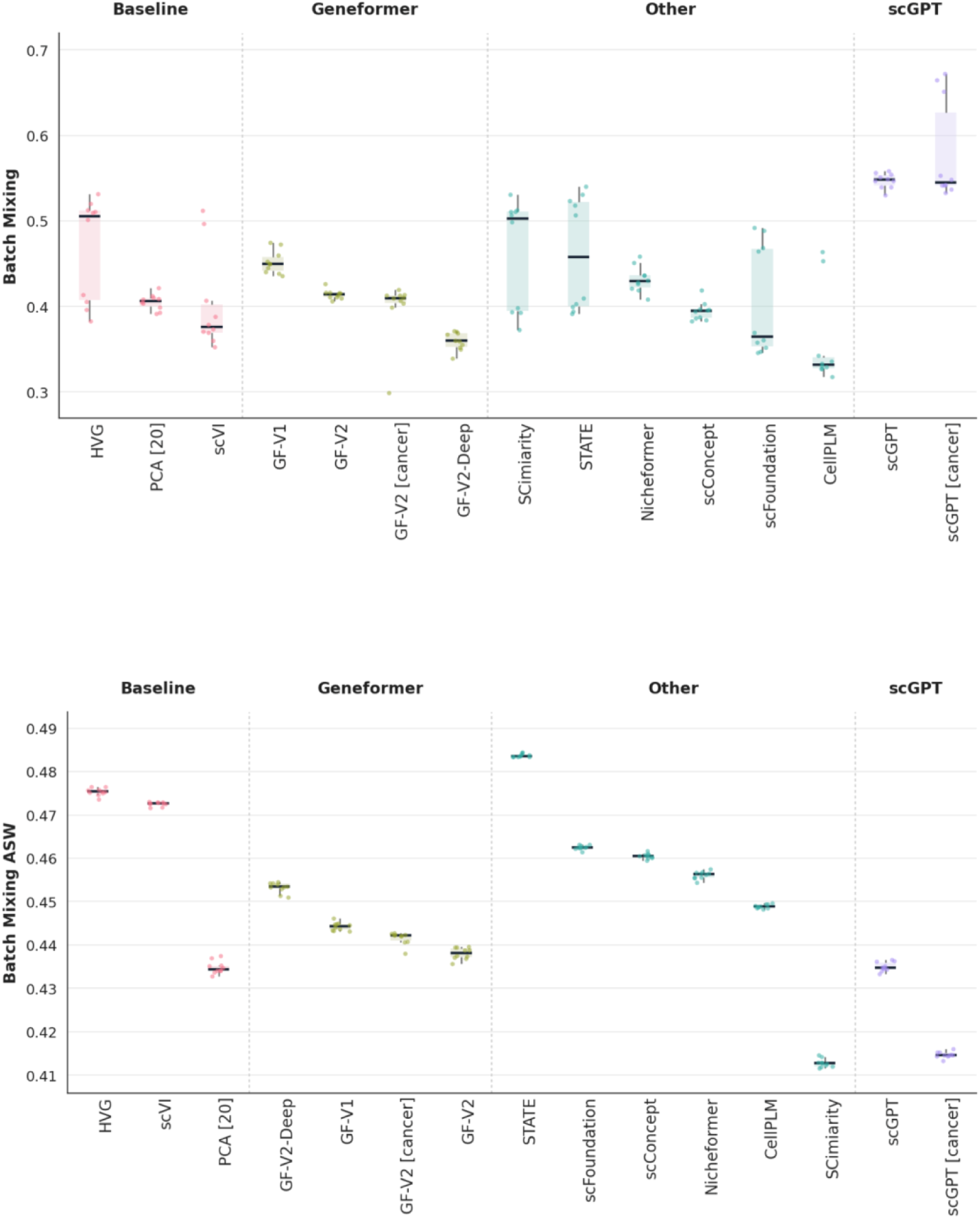
Batch mixing and batch silhouette width. **Top:** The k-nearest-neighbor batch-effect test (kBET) measures the agreement between local and global batch compositions; higher scores indicate better batch mixing. **Bottom:** Batch average silhouette width (ASW) measures separation by batch identity in embedding space using Euclidean distances; lower values indicate weaker batch separation and better mixing. Scores were calculated across 10 stratified subsamples of 10,000 cells drawn without replacement. Fifteen embedding configurations are grouped by model family and ordered within each group by median score. Boxes show the interquartile range, central lines show medians, and points represent individual subsampling runs.

**Supplementary Figure 7.**
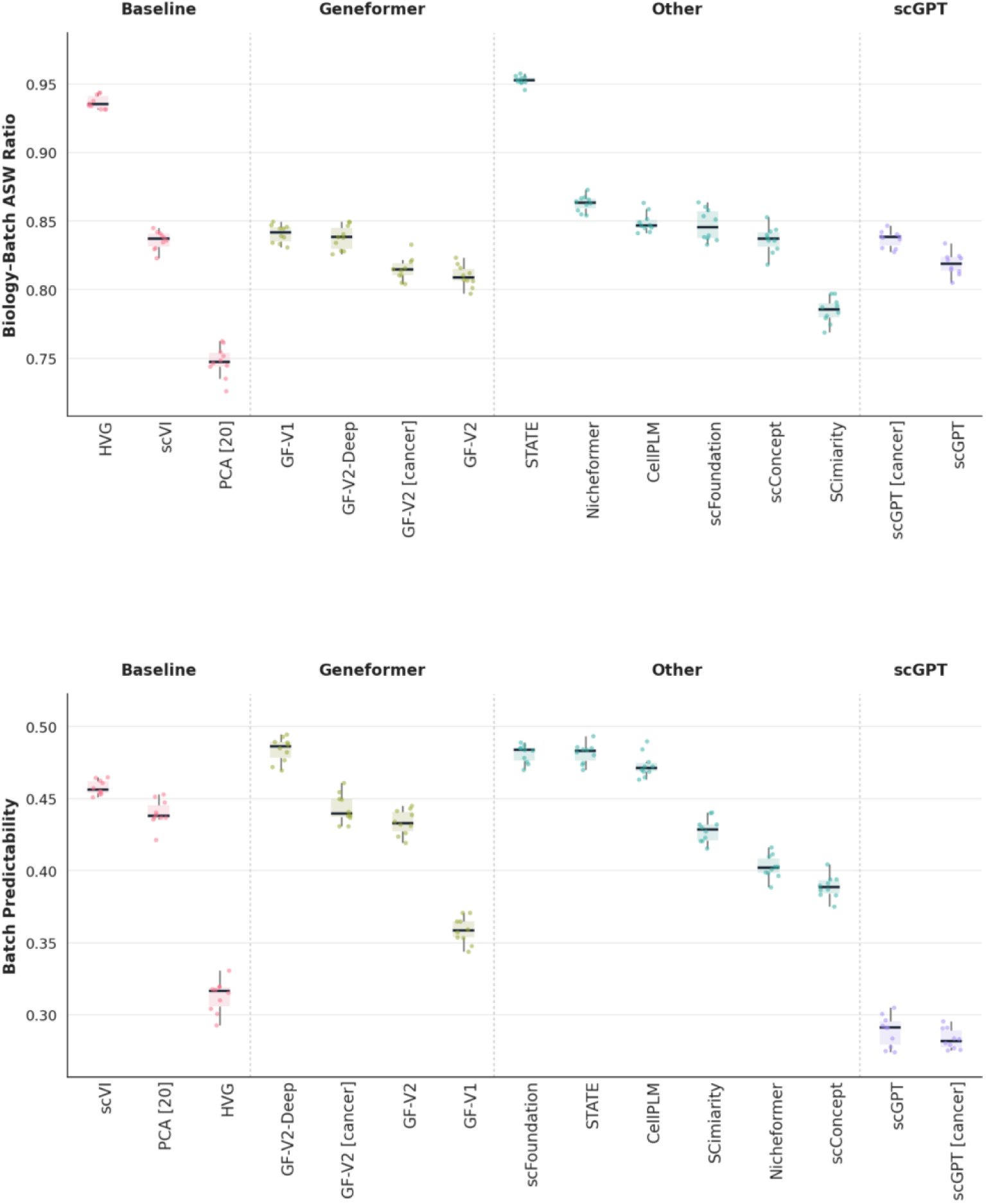
Biology-aware batch mixing and batch predictability. **Top:** The biology– batch ASW score measures batch mixing within each annotated cell type using the scaled batch silhouette width; higher values indicate better batch overlap while accounting for biological identity. **Bottom:** Batch predictability is the accuracy of a distance-weighted 15-nearest-neighbor classifier trained on 80% of cells and evaluated on a stratified 20% holdout set; lower accuracy indicates weaker residual batch effects. Scores were calculated across 10 stratified subsamples of 10,000 cells drawn without replacement. Embedding configurations are grouped by model family and ordered within each group by median score. Boxes show the interquartile range, central lines show medians, and points represent individual subsampling runs

**Supplementary Figure 8.**
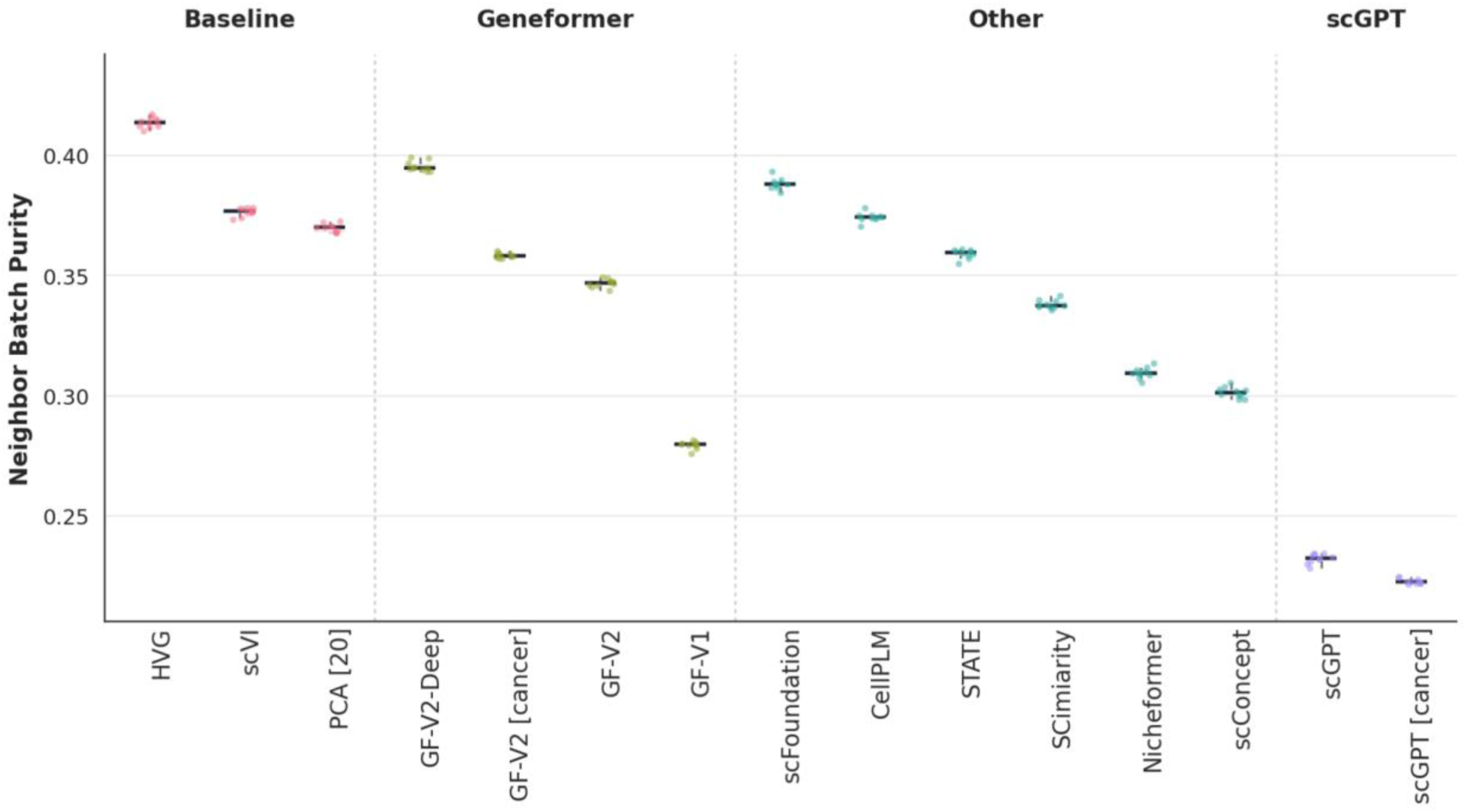
Neighbor batch purity. Neighbor batch purity is the mean proportion of each cell’s 15 nearest neighbors that share its batch label. Lower values indicate greater local batch mixing and weaker residual batch effects. Scores were calculated across 10 stratified subsamples of 10,000 cells drawn without replacement. Fifteen embedding configurations are grouped by model family and ordered within each group by median score. Boxes show the interquartile range, central lines show medians, and points represent individual subsampling runs.

**Supplementary Figure 9.**
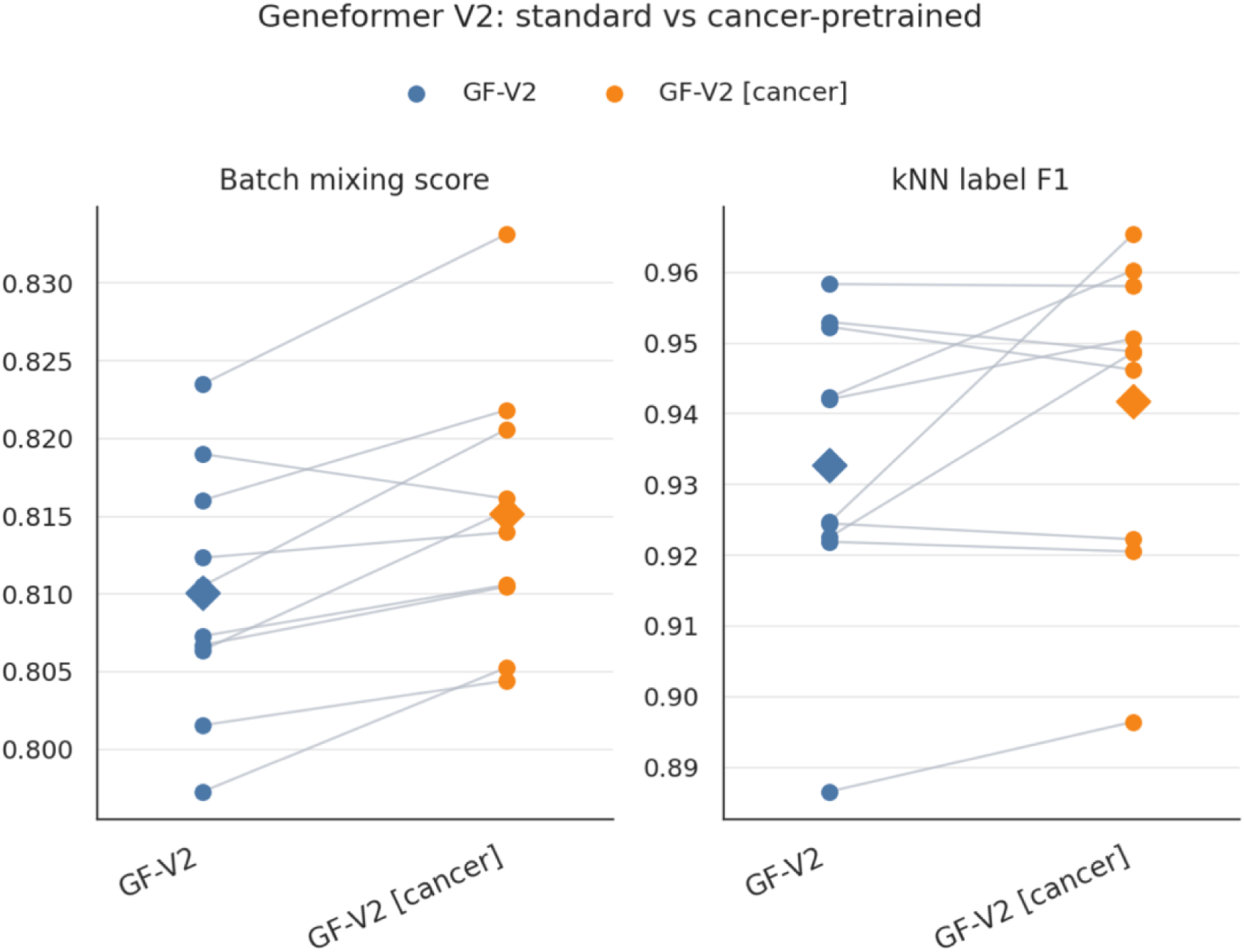
Effect of cancer-specific pretraining. Paired slope plots compare the standard Geneformer GF-V2 model with the cancer-pretrained GF-V2 [cancer] model. Cancer-specific pretraining improved both evaluated metrics. The improvement in batch mixing was significant (Wilcoxon signed-rank test, *P* = 0.006), whereas the improvement in kNN label F1 was not significant (*P* = 0.23).

**Supplementary Figure 10.**
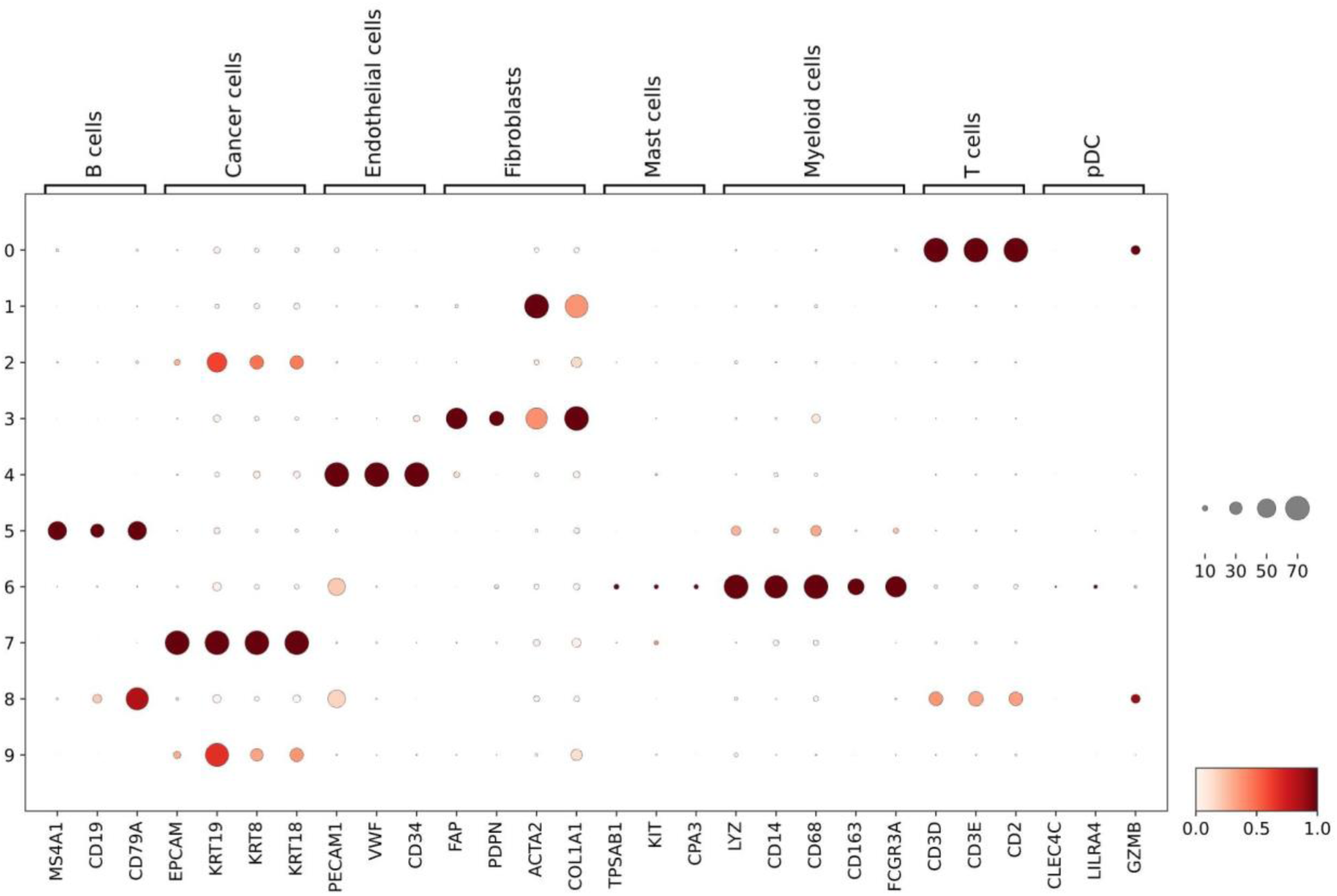
Dot plot of canonical tumor microenvironment (TME) markers. Dot plot of canonical tumor microenvironment (TME) marker genes across Leiden clusters (resolution 0.3) derived from Geneformer V1 continue embeddings in pre-treatment BRCA tumor cells (n=26,697 cells, 10 clusters). Dot size indicates the fraction of cells with detectable expression; color indicates mean log-normalized expression (scaled per gene). Embedding-derived clusters recovered expected marker structure, with the strongest enrichments including Cancer cells markers in cluster 7 (91% expressing cells), endothelial cell markers in cluster 4 (79% expressing cells), T cells markers in cluster 0 (79% expressing cells).

**Supplementary Figure 11.**
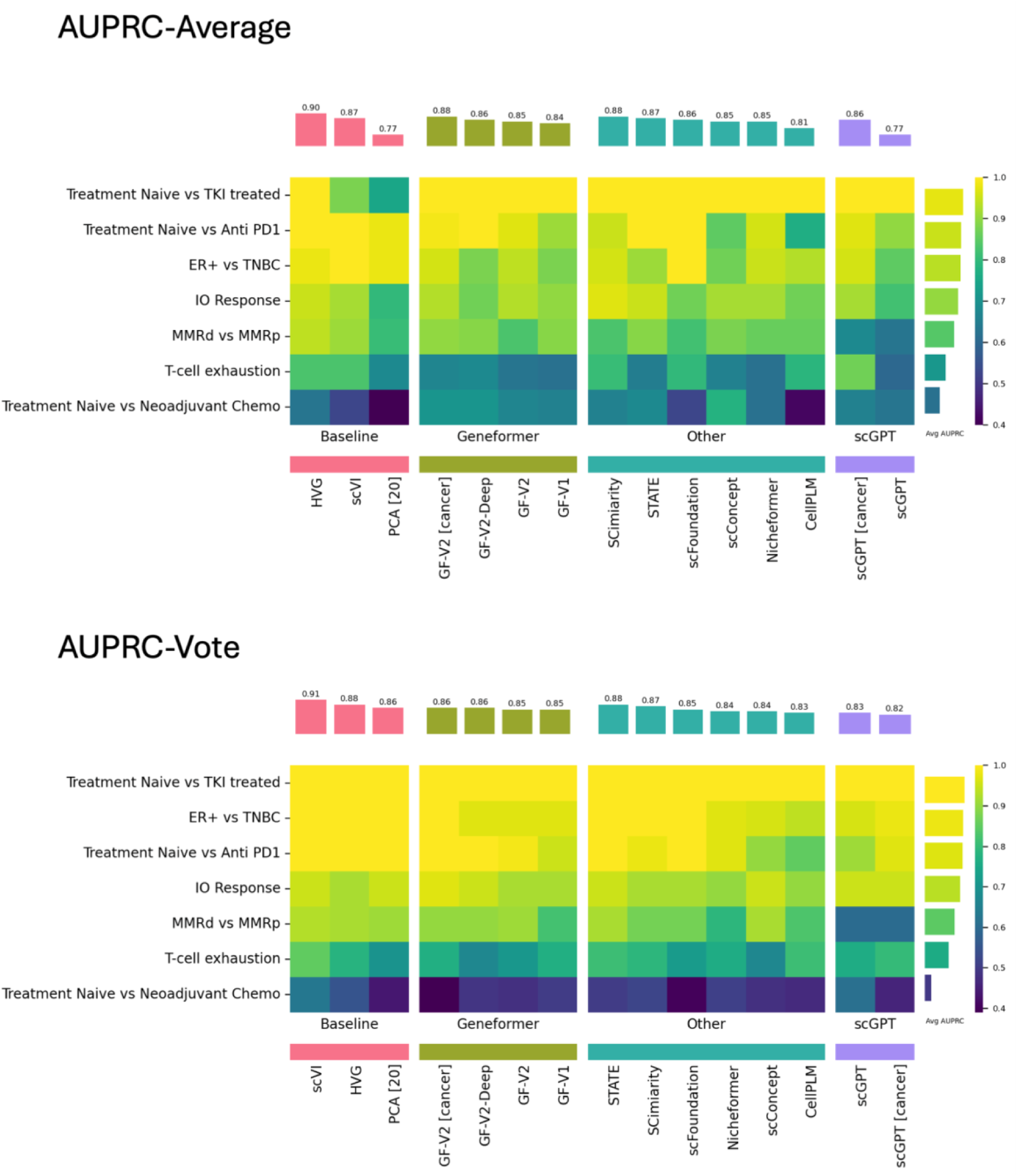
Model performance in predicting cancer-related outcomes. Heat maps compare the performance of baseline and foundation-model embeddings across seven cancer-related prediction tasks using embedding averaging (top) and majority-vote aggregation (bottom). Performance was measured by the area under the precision–recall curve (AUPRC), with higher values indicating better prediction. Each heat-map cell represents the mean AUPRC across five cross-validation folds. Bars above the heat maps show mean performance across tasks for each model, whereas bars on the right show mean performance across models for each task.

**Supplementary Figure 12.**
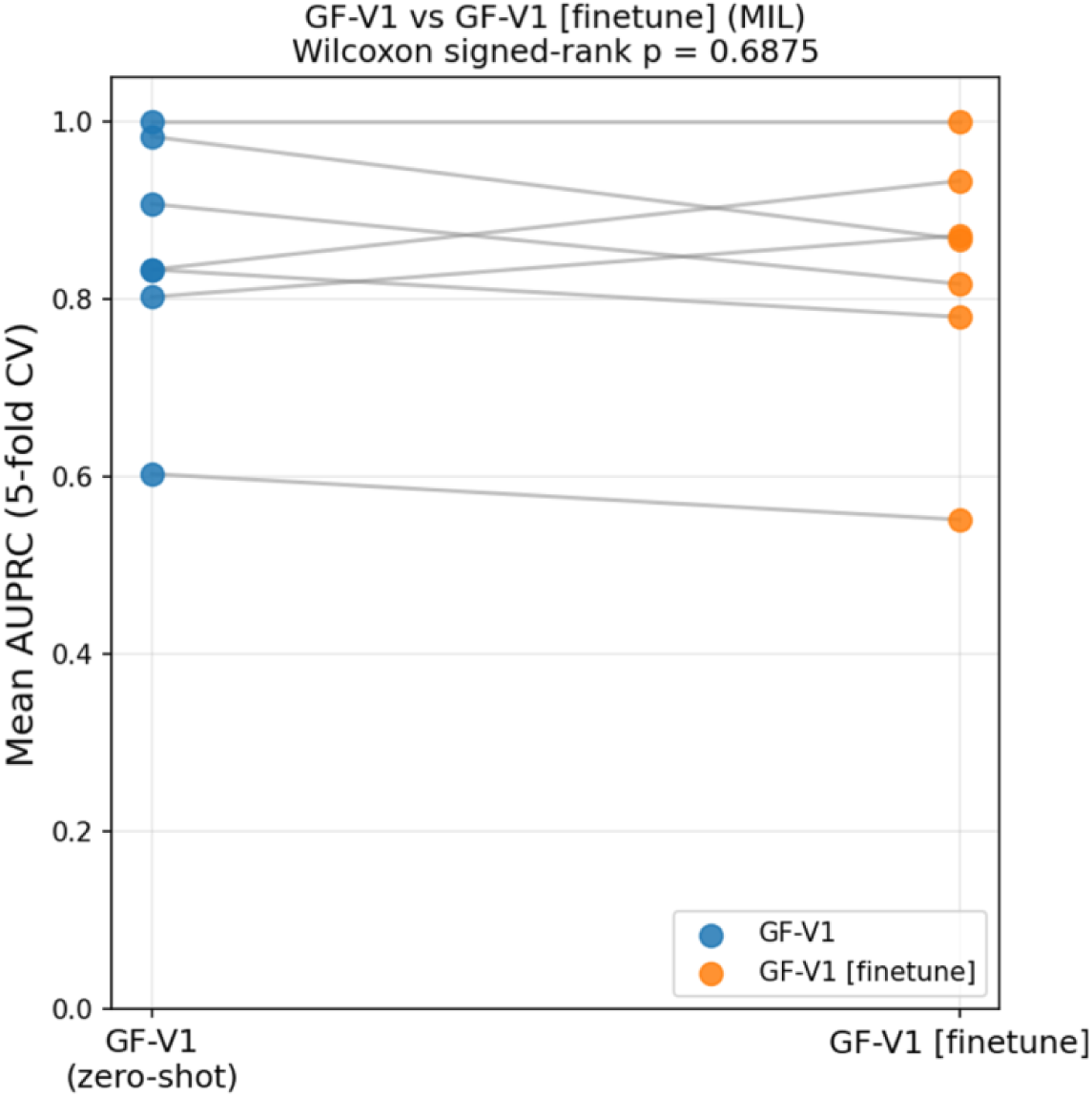
Comparison of zero-shot and fine-tuned Geneformer V1 performance. Paired comparison of zero-shot and fine-tuned Geneformer V1 (10M) across seven cancer classification benchmarks using multiple-instance learning (MIL). Points show the mean AUPRC across five cross-validation folds for each task, with lines connecting results from the same task. Fine-tuning did not significantly improve overall performance across tasks (two-sided Wilcoxon signed-rank test, *P* = 0.6875).

**Supplementary Figure 13.**
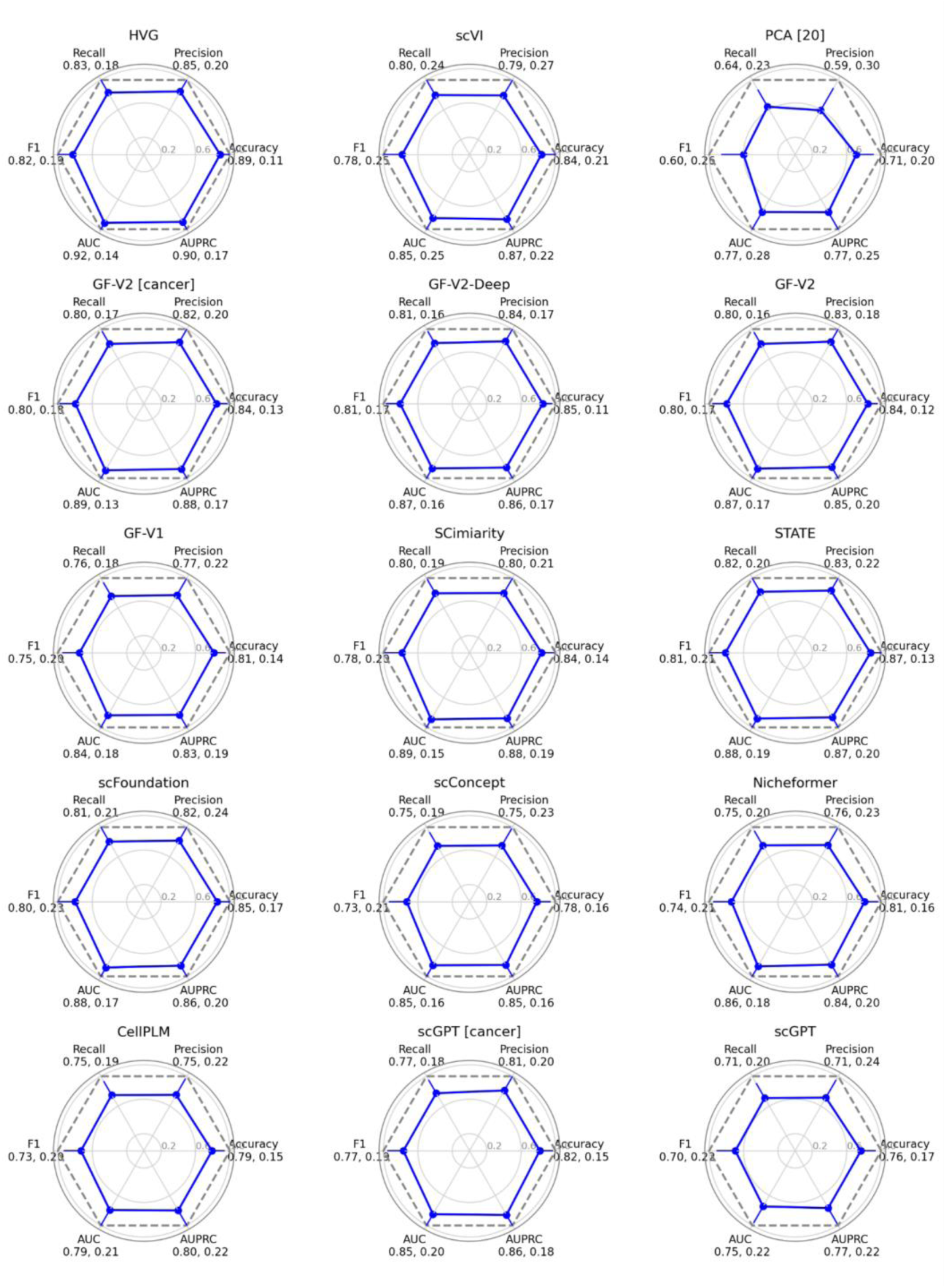
Overall classification performance using mean embedding aggregation. Radar charts summarize the performance of 15 embedding models across the cancer classification tasks. Charts are arranged in a 5 × 3 grid, with six axes representing, clockwise from the right, accuracy, AUPRC, AUC, F1 score, recall, and precision. Blue polygons connect the mean values across tasks, and outward error bars represent one standard deviation. The dashed gray polygon denotes the ideal score of 1 for all metrics. The mean and standard deviation for each metric are reported beside the corresponding axis label.

**Supplementary Figure 14.**
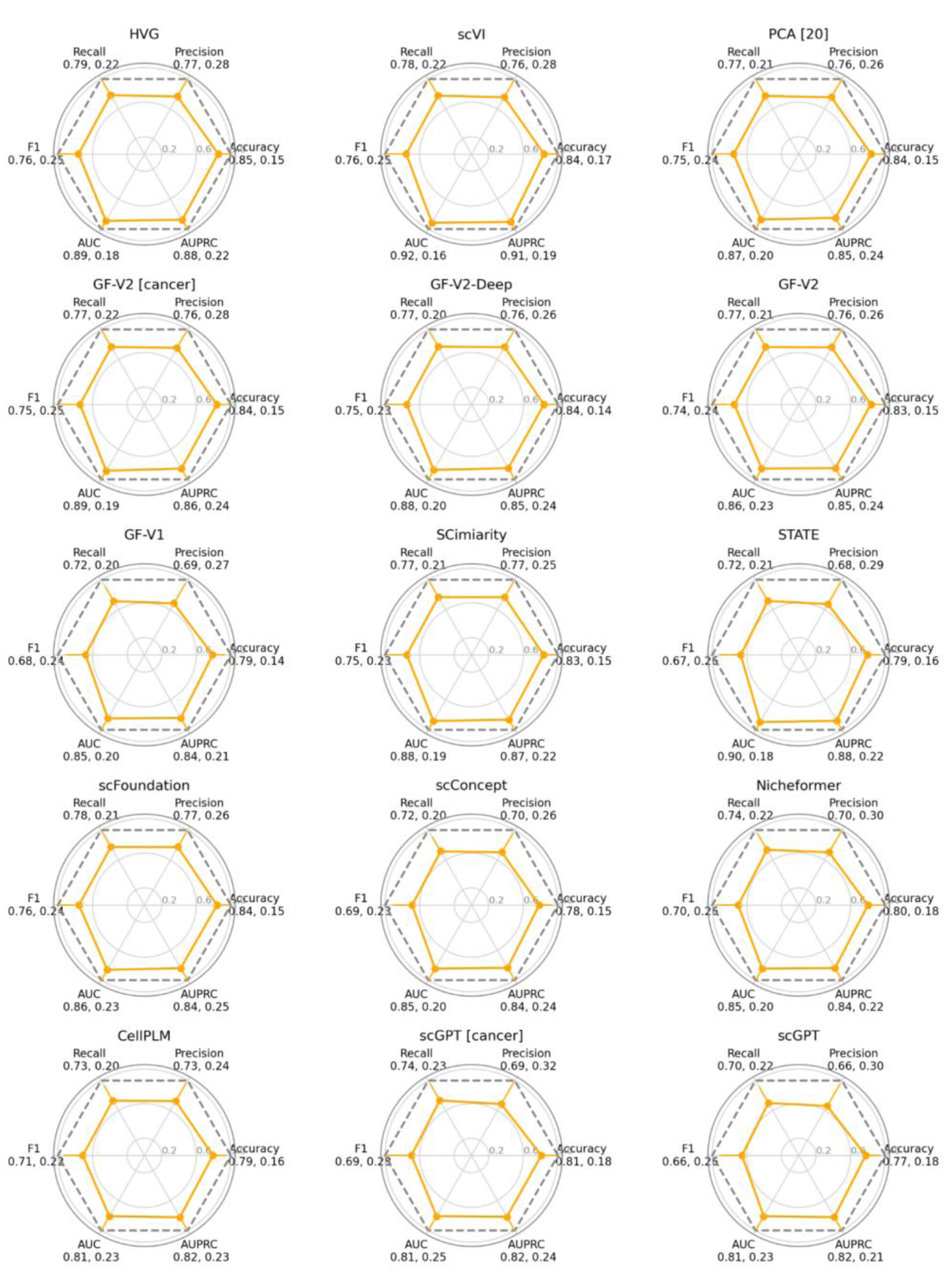
Overall classification performance using voting aggregation. Radar charts summarize the performance of 15 embedding models across the cancer classification tasks. Charts are arranged in a 5 × 3 grid, with six axes representing, clockwise from the right, accuracy, AUPRC, AUC, F1 score, recall, and precision. Blue polygons connect the mean values across tasks, and outward error bars represent one standard deviation. The dashed gray polygon denotes the ideal score of 1 for all metrics. The mean and standard deviation for each metric are reported beside the corresponding axis label.

**Supplementary Figure 15.**
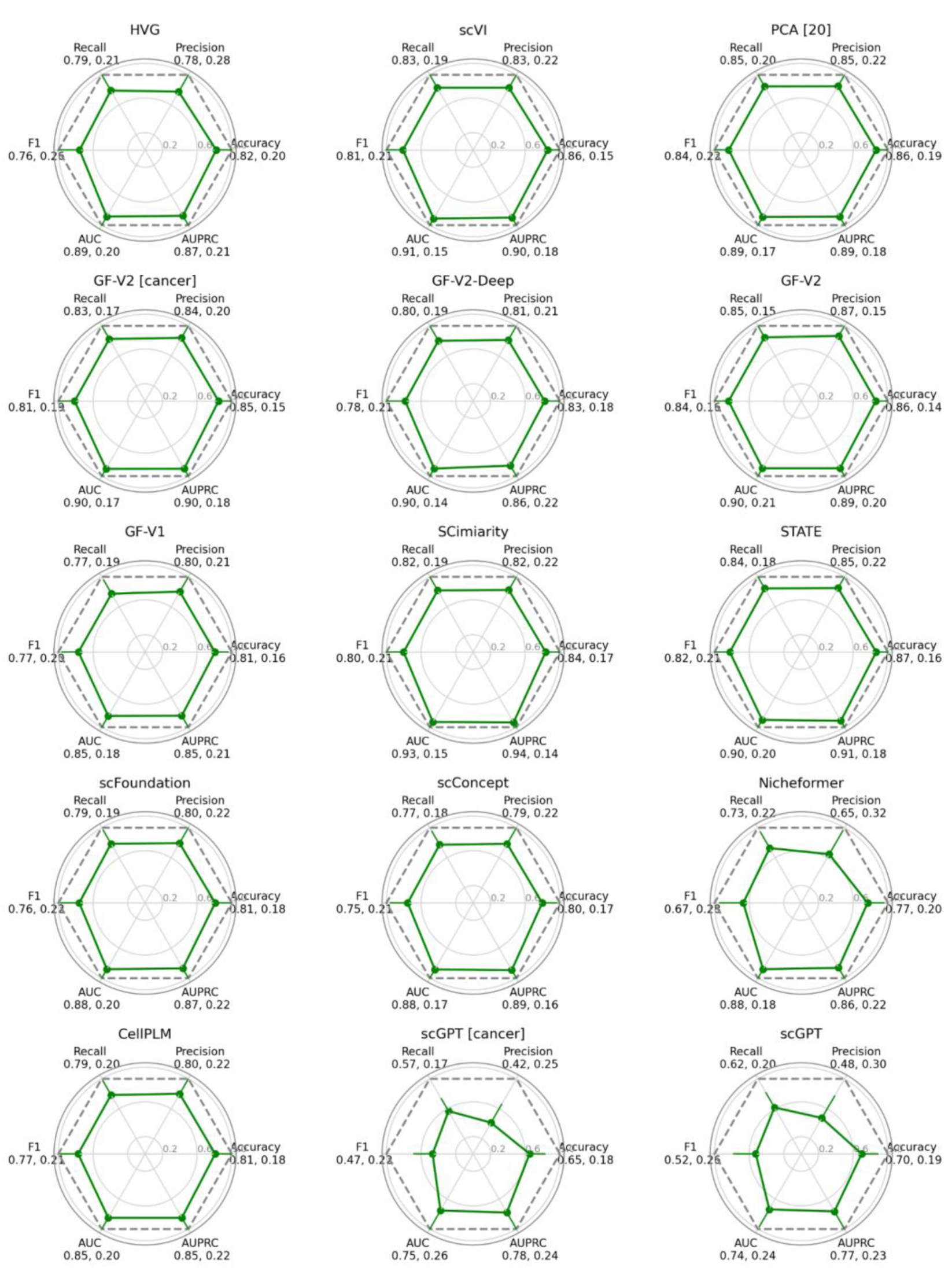
Overall classification performance using multiinstance learning (MIL) aggregation. Radar charts summarize the performance of 15 embedding models across the cancer classification tasks. Charts are arranged in a 5 × 3 grid, with six axes representing, clockwise from the right, accuracy, AUPRC, AUC, F1 score, recall, and precision. Blue polygons connect the mean values across tasks, and outward error bars represent one standard deviation. The dashed gray polygon denotes the ideal score of 1 for all metrics. The mean and standard deviation for each metric are reported beside the corresponding axis label.

## Notes

https://github.com/marakeby/scFM_eval

https://marakeby.github.io/scfm-benchscope/

### Summary of Updates

This revision expanded the number of evaluated models, provided new insights, and reported the development of agentic framework to discover and evaluate emerging scFMs

